# amica: an interactive and user-friendly web-platform for the analysis of proteomics data

**DOI:** 10.1101/2021.11.23.466958

**Authors:** Sebastian Didusch, Moritz Madern, Markus Hartl, Manuela Baccarini

## Abstract

**Summary:** Quantitative proteomics has become an increasingly prominent tool in the study of life sciences. A substantial hurdle for many biologists are, however, the intricacies involved in the associated high throughput data analysis. In order to facilitate this task for users with limited background knowledge, we have developed amica, a freely available open-source web-based software that accepts proteomic input files from different sources and provides quality control, differential expression, biological network and over-representation analysis on the basis of minimal user input. Scientists can use amica interactively to compare proteins across multiple groups, create customized output graphics, and ultimately export the results in a tab-separated format that can be shared with collaborators.

**Availability and Implementation:** The code for the application, input data and documentation can be accessed online at https://github.com/tbaccata/amica and is also incorporated in the web application. A freely available version of amica is available at https://bioapps.maxperutzlabs.ac.at/app/amica.

## Introduction

Mass spectrometry (MS)-based proteomics enables deep qualitative and quantitative characterization of any organism’s proteome which is crucial for the understanding of the underlying cell biology, physiology and biochemistry. The constant technological advancement of instruments and data-acquisition techniques, as well as the parallel development of a broad variety of methods such as proximity-dependent labeling for the study of weak or transient proteinprotein interactions (PPIs), has expanded the scope and relevance of MS-based approaches in tackling specific biological questions. As a result, proteomic approaches have become increasingly popular, but the complex analysis of MS-based proteomics data is an obstacle for many novices in the field, complicating and delaying the interpretation of experimental outcomes.

Moreover, the many software platforms available for processing proteomics raw data and their different output formats require advanced knowledge for obtaining final interpretable resuits. Several attempts to solve this problem have been made in the past. The software Perseus,^1^ for example, is widely used because it provides a graphical user interface and has extensive options for the analysis of both label-based and label-free methods. Other software tools, for example MSstats^2^ or MSnbase^3^ revolve around the R programming language. These tools have the advantage of automating many of the data processing steps that would have to be performed manually in the Perseus interface, but require knowledge of the programming language R.

Recently developed applications integrate their software into R-Shiny apps, allowing for interactive user interactions and visualizations of the output from the MaxQuant package, one of the most widely used software platforms in MS-based proteomics.^4^ LFQ-Analyst^5^ enables the automatic analysis of label-free data, Eatomics^6^ allows for the input of enhanced experimental designs and ProVision^7^ can process label-free and TMT labeled data and integrates PPI networks. These tools provide appealing solutions for first-time users but are limited to output from MaxQuant. In addition, none of these tools allow the systematic comparison of proteins across multiple groups or the integration and comparison of multiple proteomics experiments.

As a solution to these issues, we have developed amica, a user-friendly web-based platform for comprehensive quantitative proteomics data analysis that can automatically handle multiple database search tool outputs such as for instance data from FragPipe - a recent but increasingly popular open-source software package^8,9^ - as well as any generic tab-separated dataset such as RNA-seq data and datasets that were previously analyzed with amica. By allowing the upload of a second amica file to be compared with the current data input, the tool enables multi-omics integration.

## Methods

### Example data set

An example data set was selected from PRIDE identifier PXD016455.^10^ This includes raw files for four groups from an interaction proteomics study focusing on PGRMC1, a protein from the MAPR family with a variety of cellular functions. In this study, MIA PaCa-2 cells were stably transfected with a PGRMC1-HA plasmid and Co-IPs of PGRMC1 interacting proteins were isolated from cells expressing PGRMC1-HA, as well as from non-transfected parental MIA PaCa-2 cells as a negative control, with and without AG-205 treatment (a PGRMC1-specific inhibitor).

### Data analysis

MaxQuant (version 1.6.17.0) was used to analyze the raw files. As search database, UniProt UP000005640 (downloaded on 10th September 2021) was used, with Trypsin/P as proteolytic enzyme allowing for two missed cleavages. The match between runs (MBR) feature was not used. Oxidation on methionine and protein N-terminal acetylation were set as variable modifications, and Carbamidomethylation of Cystein was set as fixed modification. Label-free quantification and normalization was performed with the MaxLFQ algorithm.

Additionally, FragPipe (version 16) with MS-Fragger^11^ (version 4.0.0) and Philosopher^9^ (version 4.0.0) was used to analyze the raw files. Label-free quantification and normalization was performed with the MaxLFQ algorithm by IonQuant^12^ (version 1.7.5). The same search database as well as variable and fixed modifications as for MaxQuant were used for FragPipe. Peptide validation was executed by Percolator^13^ (version 3.05). The MBR feature was not used.

The output from MaxQuant and FragPipe was analyzed using the same analysis parameters in amica. Briefly, proteins with at least 2 Razor + unique peptides, at least 3 MS / MS counts, and valid values in 3 out of 5 replicates in at least one group were considered quantified. LFQ intensities of quantified proteins were log2-transformed and missing values were imputed from a normal distribution downshifted 1.8 standard deviations from the mean with a width of 0.3 standard deviations. Differential expression analysis was performed with limma.^14^

## Results and discussion

amica was developed as a user-friendly R-Shiny app with interactive and customizable visualizations that can be exported in a publication-ready vector graphic format.

amica requires three input files: a) a database search result such as MaxQuant’s proteinGroups.txt file, FragPipe’s combined_proteins.txt file or a custom tab-separated file b) a tab-separated file denoting the experimental design in the experiment, i.e. the mapping of samples to distinct biological groups and c) a tab-separated file containing a contrast matrix that specifies the desired pairwise group comparisons to be made. The parameters for the analysis address filtering by valid values, normalization, imputation of missing values, differential abundance testing with limma^14^ or DEqMS^15^ and can be changed in the user interface. The results of the analysis can be downloaded in a custom amica format, which can later be used as the input file for subsequent re-inspection in amica. amica’s landing page displays online documentation, a link a user manual and a link to download the formatted example data sets (Fig. 1).

**Figure 1:**
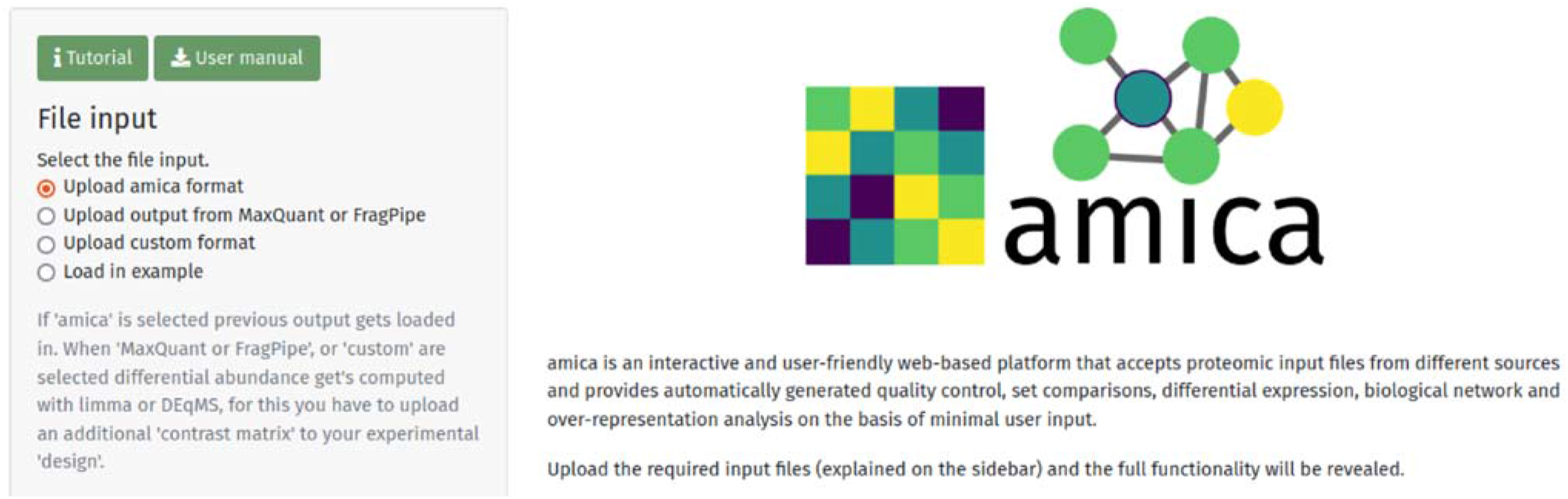
amica’s user interface visible after uploading the required input files.

Quality control can be visualized for raw intensities, LFQ intensities, normalized and imputed intensities. This includes barplots for the number of identified proteins (Fig. 2a), contaminants, missing values, most abundant proteins per sample; as well as density plots, boxplots for intensities and coefficent of variations, correlation plots (Fig. 2b), scatter plots (Fig. 2c), and a principal component analysis (PCA) plot (Fig. 2d).

**Figure 2:**
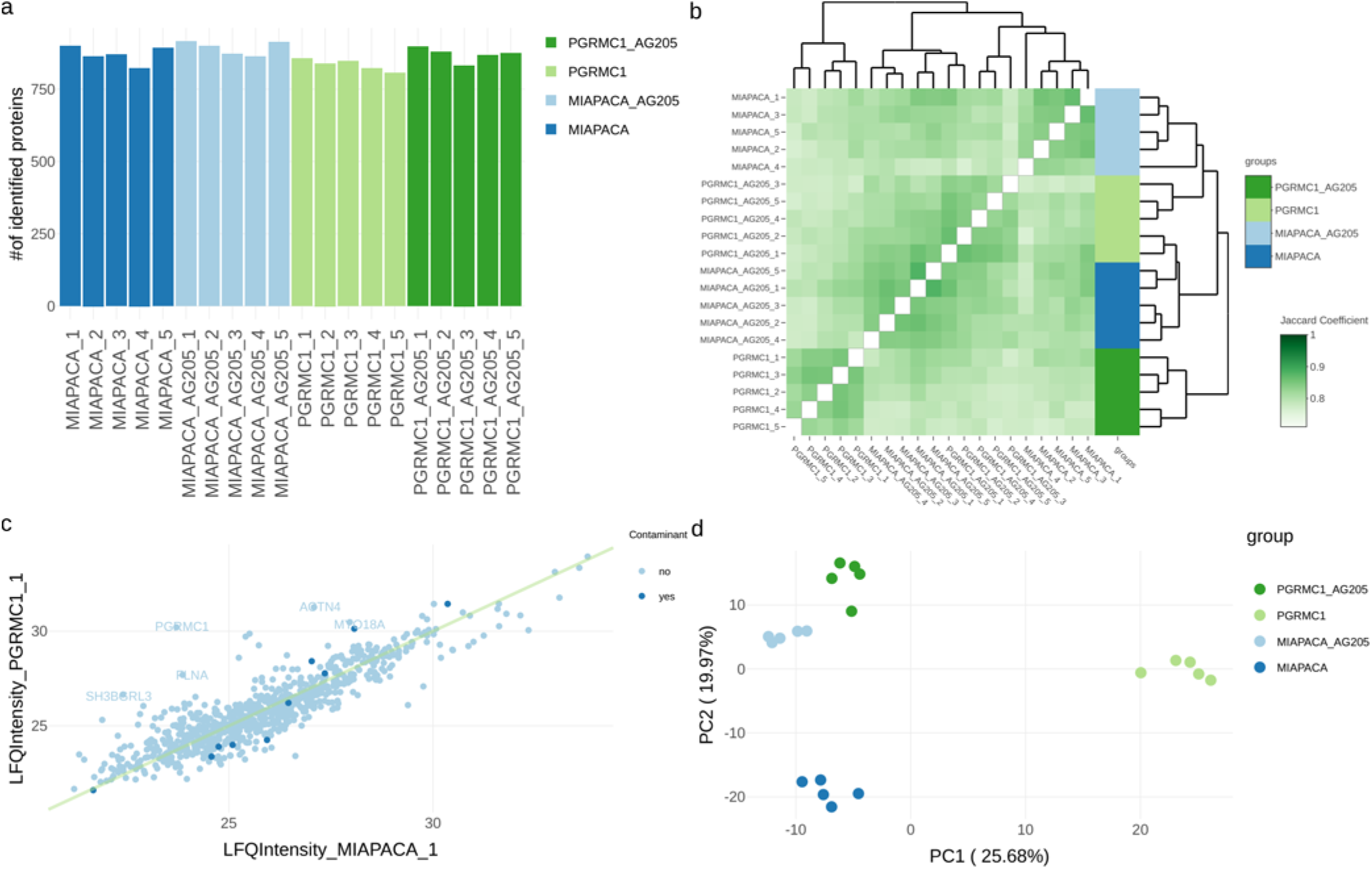
Examples of amica graphics, produced with the provided example dataset^10^. (a) The number of identified proteins can be visualized in a bar plot. (b) The overlap of identified proteins between samples is shown in an interactive heatmap.^16^ Users have the choice of displaying the overlap as the number of shared proteins, Overlap coefficient or Jaccard coefficient. (c) Scatter plots can be plotted using different intensities and samples, allowing users to interactively highlight proteins. User can choose drawing a straight line, a line from a linear regression or no line in the scatter plot. (d) PCA plot.

Differentially abundant proteins can be visualized as volcano (Fig. 3a) - and MA - plots for single group comparisons. Unlike other R-Shiny apps for proteomics data analysis, amica can generate UpSet plots^17^ (Fig. 3b) and Euler diagrams (Fig. 3c) for visualizing the overlap of significant proteins from multiple selected group comparisons. Statistically significant proteins are displayed in a data table and can be further filtered, allowing for the immediate inspection of relevant proteins specific to a comparison in subsequent visualizations. These include foldchange plots (Fig. 3d), profile plots, interactive heatmaps and Dot plots (Fig. 3e).

**Figure 3:**
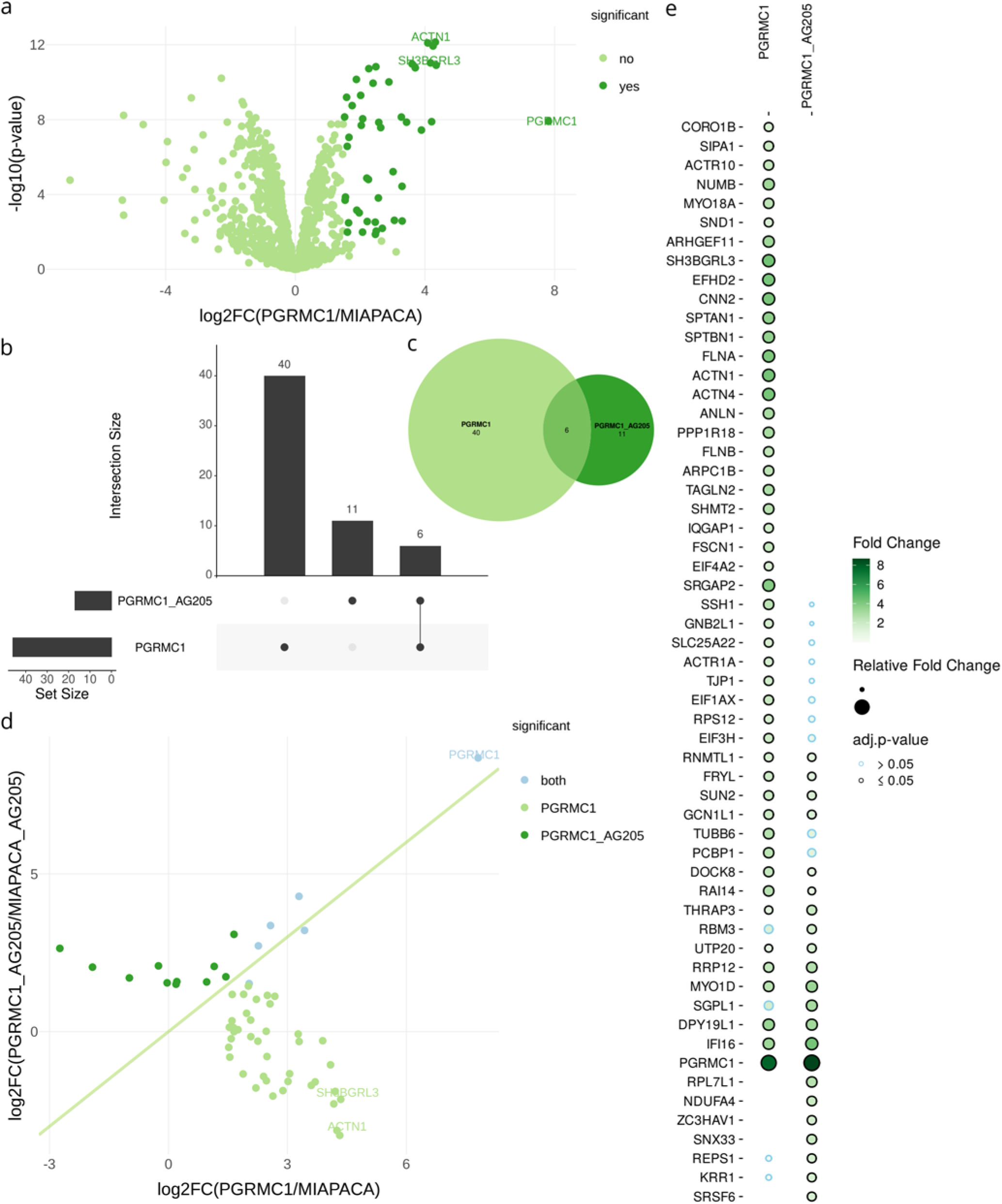
(a) Differentially abundant proteins can be represented with volcano plots. Proteins specific to group comparisons are visualized in Upset plots (b), Euler diagrams (c), fold-change plots (d), or dot plots (e).

For human proteomics studies, amica allows users to map proteins to a PPI network from the IntAct database^18^ (Fig. 4a), and enables the retrieval of subcellular localization predictions from the humancellmap database^19^. As an example, selecting a particular cellular compartment in the interactive web interface will highlight all proteins that map to this localization. This information can be downloaded in gml format for visualization in a network visualization tool such as Cytoscape^20^. A functional enrichment analysis of differentially abundant proteins using gprofiler2^21^ assists in building hypotheses on the underlying biology (Fig. 4b).

**Figure 4:**
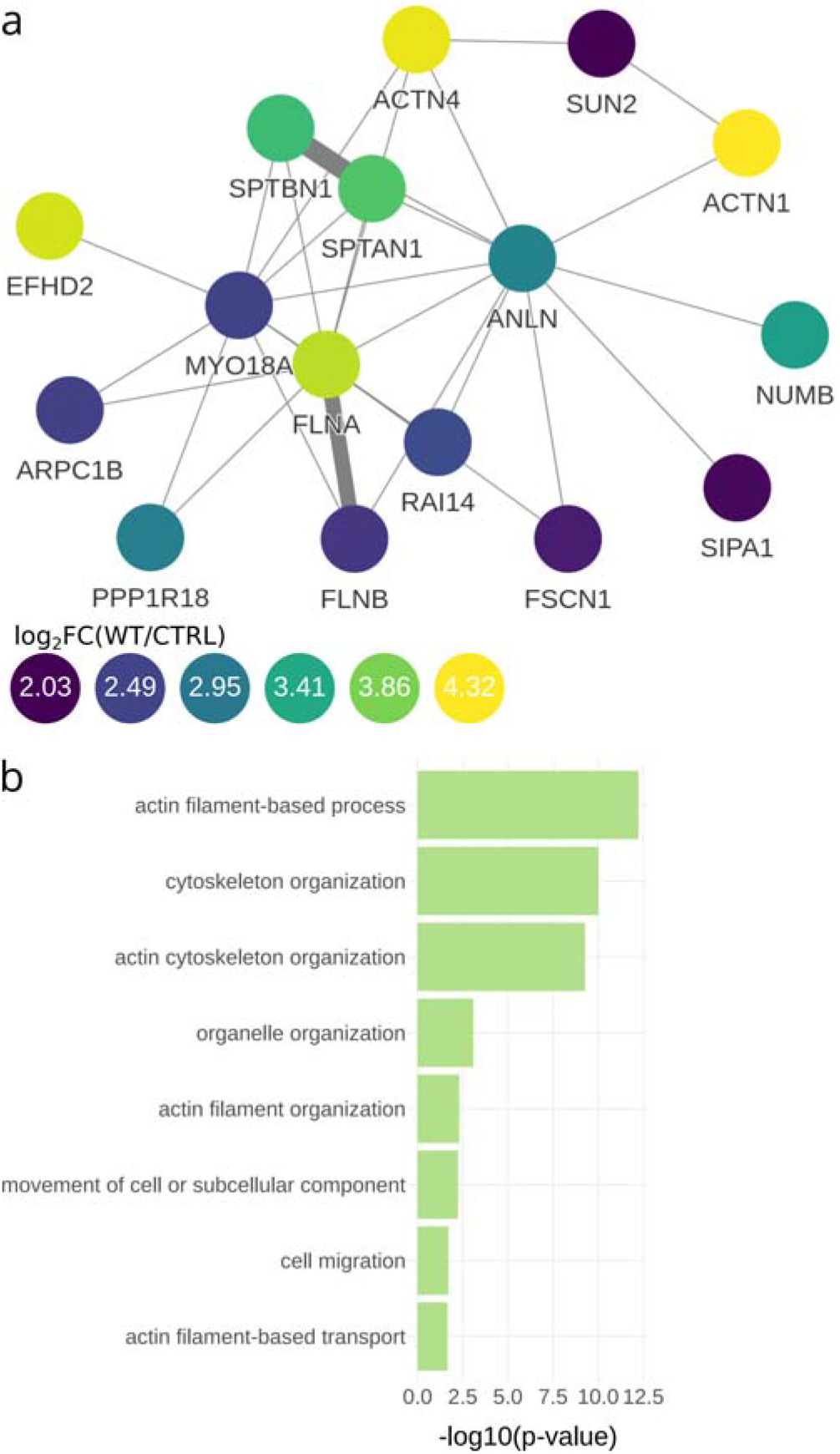
(a) Differential abundant proteins can be visualized in a PPI-Network from the IntAct database. (b) Results from a functional enrichment analysis using gprofiler2.

Finally, amica accepts and analyzes any generic tab-separated data set, provided a specification file is uploaded that maps the relevant search engine specific columns to a standard format. This feature, combined with the possibility to upload previously analyzed datasets to be compared with the current data input, facilitates multi-omics integration.

## Conclusions

We have developed amica as a general software tool for analyzing and visualizing MS-based proteomics data. amica’s user-friendly interface provides a customizable data analysis workflow, and the results of the analysis can be conveniently exported, shared and reloaded into the amica environment for re-inspection at a later time. The data analysis workflow in amica includes quality control and standard differential expression testing, as well as the integration of PPI networks and pathway and gene ontology enrichment analysis for differentially abundant proteins - all of which can help researchers to better interpret the biology underlying the results of their proteomics experiment. The code for the application and online documentation can be found at https://github.com/tbaccata/amica and the software is available at https://bioapps.maxperutzlabs.ac.at/app/amica.

## Supporting information

amica Manual

## Acknowledgement

The authors thank the Baccarini lab, the Max Perutz Labs MS facility, Jörg Menche and his group, Matthias Bärnthaler and the IT team for feedback and helpful comments.

This work was supported by Austrian Research Fund (W1261 to MB, SFB F70, F7007 to MH). The authors declare no competing financial interest.

## Supporting Information Available

The following files are available free of charge.

- Supplementary Note 1: Link to a user manual demonstrating every available function and output graphics, as well as extensive tutorials.

