## Supplementary material for "amica: an interactive and user-friendly web-platform for the analysis of proteomics data": amica Manual

April 7, 2022

**8 Comparison with other R-Shiny apps for proteomics data analysis** **41**

### Chapter 1

### Introduction

amica is a user-friendly and interactive web-based platform for the analysis and visualization of proteomics data. amica accepts proteomic input files from different sources and provides quality control, differential expression, biological network and over-representation analysis on the basis of minimal user input. Scientists can use amica interactively to compare proteins across multiple groups, create customized output graphics, and ultimately export the results in a tab-separated format that can be shared with collaborators. Examples of amica graphics, produced with the provided example dataset [1] are shown in Figure 1.

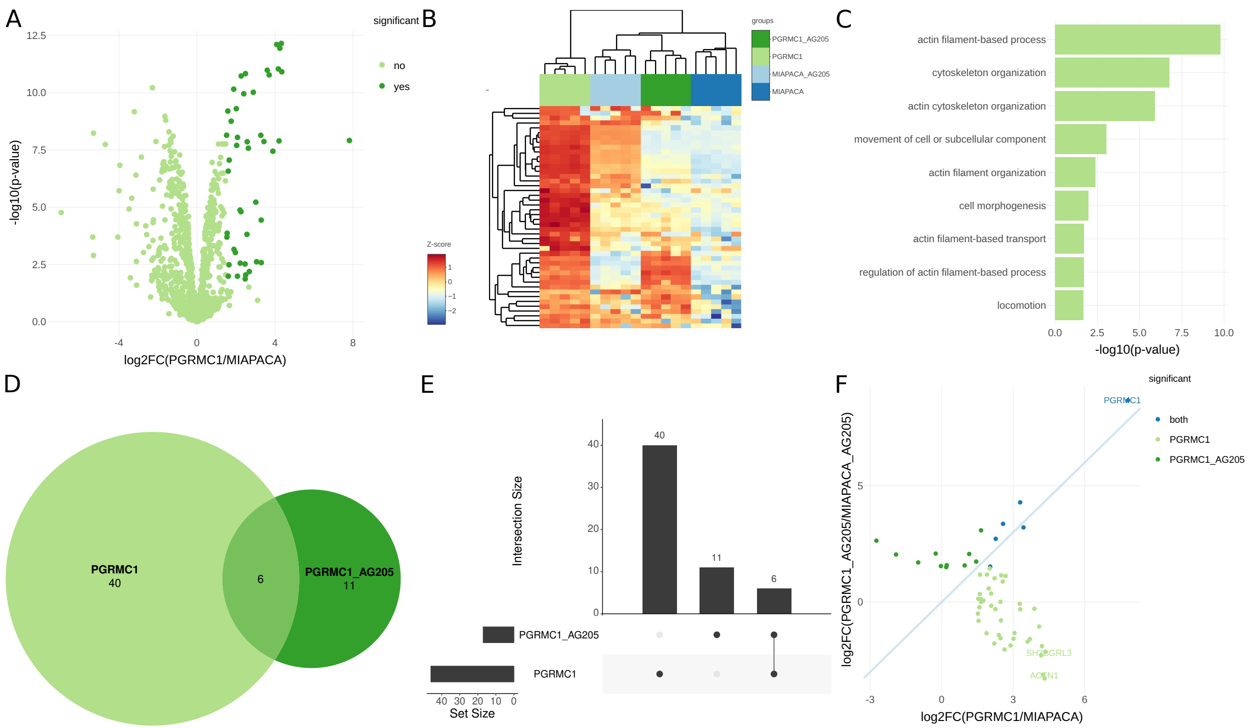

Figure 1: Differentially abundant proteins can be represented with volcano plots (A) or interactive heatmaps [2] in (B). Results from functional enrichment in (C). Proteins specific to group comparisons are visualized in Euler diagrams (D), Upset plots (E) or fold-change plots (F).

### Chapter 2

### User input

#### 2.1 Quick Start

- Go to https://bioapps.maxperutzlabs.ac.at/app/amica.
- Choose one out of 4 upload options (Fig. 2).
  - upload amica file and an experimental design.
  - upload MaxQuant's [3] proteinGroups.txt or FragPipe's [4, 5] combined_proteins.tsv together with an experimental design and a contrast matrix.
  - upload a custom tab-separated file with a specification file, an experimental design and a contrast matrix.
  - Select the "load in example" option.
- Click on "Upload" (and on "Analyze" if you have selected MaxQuant, FragPipe or custom input).
- In the main tab menu the tabs QC, Differential abundance and Compare amica experiments become visible.

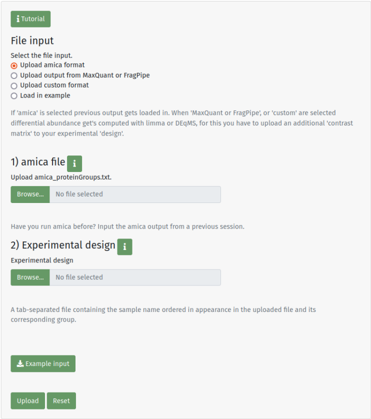

Figure 2: Upload options.

#### 2.2 Accepted input formats

amica can read in common database search tool output formats, custom formats and its own tab-separated format. It is able to achieve this by mapping file specific column names into common features present in proteomics data. These include a unique protein id, a gene name, intensities, peptide counts, spectral counts and others. This chapter describes the correct formatting of input files.

Example files for all input formats can be downloaded in the input tab of amica.

##### 2.2.1 MaxQuant's proteinGroups.txt

For MaxQuant label-free quantification (LFQ) output following columns are parsed:

| Variable name | Column name/prefix | comment |
|---|---|---|
| proteinId | "Majority protein IDs" | |
| geneName | "Gene names" | |
| intensityPrefix | "LFQ Intensity <sample>" | |
| Imputed Int. prefix | - | Get's calculated |
| abundancePrefix | "iBAQ <sample>" | |
| razorUniqueCount | "Razor + unique peptides" | specific column of summarized razor+unique count |
| razorUniquePrefix | "Razor + unique peptides <sample>" | corresponds to razor+unique count of a sample |
| spectraCount | "MS/MS count" | |
| contaminantCol | "Potential contaminant" | |

amica automatically filters out reverse hits and proteins only identified by site.

##### 2.2.2 FragPipe's combined_proteins.tsv

For FragPipe/Philosopher LFQ output following columns are parsed:

| Variable name | Column name/prefix | comment |
|---|---|---|
| Default parameters | | |
| proteinId | "Protein ID" | |
| geneName | "Gene Names" | |
| intensityPrefix | "<sample> Razor Intensity" | |
| Imputed Int. prefix | - | Get's calculated |
| abundancePrefix | - | |
| razorUniqueCount | "Unique Stripped Peptides" | |
| razorUniquePrefix | "<sample> Razor Spectral Count" | |
| spectraCount | "Summarized Razor Spectral Count" | |
| FragPipe v16 (MSFragger v3.3, Philosopher v4.0.0) | | |
| proteinId | "Protein ID" | |
| geneName | "Gene Names" | |
| intensityPrefix | "<sample> Intensity" | |
| Imputed Int. prefix | - | Get's calculated |
| abundancePrefix | - | |
| razorUniqueCount | "Combined Total Peptides" | |
| razorUniquePrefix | "<sample> Razor Spectral Count" | |
| spectraCount | "Combined Spectral Count" | |
| FragPipe v17 (MSFragger v3.4, Philosopher v4.1.0) | | |
| proteinId | "Protein ID" | |
| geneName | "Gene" | |
| intensityPrefix | "<sample> MaxLFQ Intensity" | |
| Imputed Int. prefix | - | Get's calculated |
| abundancePrefix | - | |
| razorUniqueCount | "Combined Total Peptides" | |
| razorUniquePrefix | "<sample> Razor Spectral Count" | |
| spectraCount | "Combined Spectral Count" | |

##### 2.2.3 amica format

This format can be downloaded in the input tab after completing the analysis of an uploaded dataset. It is meant to be a stable, sharable output format of an analyzed dataset and the differential expression analysis is not re-computed.

When downloading amica's tab-separated protein groups file following columns are present:

| Variable name | Column name/prefix | Mandatory | Comment |
|---|---|---|---|
| proteinId | "Majority.protein.IDs" | yes | all values need to be unique |
| geneName | "Gene.names" | yes | |
| intensityPrefix | "LFQIntensity_<sample>" or "Intensity_<sample>" | no | MQs "LFQ intensity " columns |
| Imputed Int. prefix | "ImputedIntensity_" | yes | Imputed and normalized intensities |
| abundancePrefix | "iBAQ_<sample>" | no | MQs "iBAQ " columns without "iBAQ peptides" |
| razorUniqueCount | "razorUniqueCount" | no | MQs "razor+unique count" column |
| razorUniquePrefix | "razorUniqueCount_<sample>" | no | MQs "razor+unique count" columns (per sample) |
| spectraCount | "spectraCount" | no | MQs "MS/MS count" column |
| contaminantCol | "Potential.contaminant" | no | |
| quantCol | "quantified" | no | Proteins that have been quantified |
| pvalPrefix | "P.Value_" | no | e.g "P.Value_group1__vs__group2" |
| padjPrefix | "adj.P.Val_" | no | e.g "adj.P.Val_group1__vs__group2" |
| logfcPrefix | "logFC_" | yes | e.g "logFC_group1__vs__group2" |
| avgExprPrefix | "AveExpr_" | no | e.g "AveExpr_group1__vs__group2" |
| comparisonInfix | "__vs__" | yes | e.g "logFC_group1__vs__group2" |

###### Additonal information

- Only mandatory columns (unique protein id, gene name, imputed intensities and at least one logfcPrefix column) need to be present in the data for amica to read it in (see section 7.1).
- IntensityPrefix, ImputedIntensityPrefix and abundancePrefix columns are log2 transformed, all 0s need to be converted to NAs. No INF values are allowed.
- ImputedIntensityPrefix columns should only contain filtered, imputed and normalized values.
- quantCol: All proteins passing spectraCount, razorUniqueCount and filtering on valid values thresholds (see section 2.4) that have been quantified are set to "+" in this column. Otherwise no value ("") is written in the column. If no `quantified` column is provided complete cases (data containing no missing values) of all `ImputedIntensity` columns and all columns containing the group comparison infix `__vs__` are set to be quantified.
- comparisonInfix: The infix is important to retrieve the group ids from a group comparison (e.g for downstream visualizations like heatmaps). The groups before and after the `__vs__` infix need to match with groups defined in the uploaded experimental design.
- razorUniqueCount is a column, razorUniquePrefix is the prefix to the count per sample, but they may very well have the same value (just like in MaxQuant's `proteinGroups.txt`)

Proteins inferred from reverse hits and peptides "only identified by site modifications" are not to be written into amica's output.

##### 2.2.4 Custom format

Users can upload a custom tab-separated file, along with an experimental design and a contrast matrix. Additionally, a **specification file** (explained in section 2.3.3) needs to be uploaded that maps relevant database search tool specific columns to amica's format.

#### 2.3 Additional files

##### 2.3.1 Experimental design

A tab-separated **experimental design** maps samples from the input file to experimental groups or conditions. An example is shown here:

| samples | groups |
|---|---|
| replicate1_WT | WT |
| replicate2_WT | WT |
| replicate3_WT | WT |
| replicate1_TRTMT | TRTMT |
| replicate2_TRTMT | TRTMT |
| replicate3_TRTMT | TRTMT |
| ... | ... |
| replicate1_KO | KO |
| replicate2_KO | KO |
| replicate3_KO | KO |

The sample names in the **samples** column need to match to the column names of the input file in the order of the input file, e.g in MaxQuants output all sample names would be prefixed by "LFQ intensity" (e.g "LFQ intensity replicate1_WT", "LFQ intensity replicate2_WT", and so on).

##### 2.3.2 Contrast matrix

A tab-separated contrast matrix needs to be uploaded when the analysis starts from scratch (from MaxQuant's `proteinGroups.txt`, FragPipe's `combined_proteins.tsv` or a custom format). An example is shown here:

| group1 | group2 |
|---|---|
| WT | KO |
| TRMT | KO |
| WT | TRMT |

The contrast matrix tells amica which group comparisons to perform. The column names of this file can be freely chosen, but column names must be provided. The comparison group1-group2 is performed for each row in this file. To change the sign of the fold changes, the position of the groups needs to be switched in the file (e.g group2-group1, change KO and WT in the example table). The groups in both columns need to be present in the experimental design file.

##### 2.3.3 Specification file

For custom file uploads, a **specification file** needs to be uploaded that maps relevant database search tool specific columns to the data. The file has two columns, Variable and Pattern:

| Variable | Pattern | Mandatory |
|---|---|---|
| proteinId | ... | yes |
| geneName | ... | yes |
| intensityPrefix | ... | yes |
| abundancePrefix | ... | no |
| razorUniqueCount | ... | no |
| razorUniquePrefix | ... | no |
| spectraCount | ... | no |
| contaminantCol | ... | no |

If `contaminantCol` is provided all protein groups that might be potential contaminants must be indicated by a "+" character just like in MaxQuant. If a non-mandatory column is not available in the custom data no Pattern should be written into the specification file.

All intensities prefixed with the intensityPrefix and abundancePrefix should not be log2-transformed, as amica log2-transforms them by default.

#### 2.4 Analysis options

After the user has successfully uploaded the required input files, the following analysis parameters can be selected (Fig. 3):

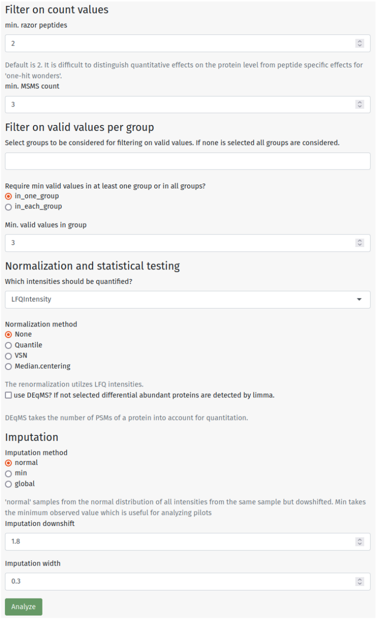

Figure 3: Upload options.

##### Filtering on minimum count values

- min. razor + unique peptide count.
- min. MS/MS count.

##### Filtering on valid values per group

- Groups can be selected for filtering on valid values → if no group is selected all groups are considered.
- Filters on valid values should be applied in at least one group or in each group.

##### Normalization and Differential expression analysis

- Intensities for normalizing and analyzing differential abundance can be chosen. Default are LFQ intensities for MaxQuant and FragPipe.
- (Re-) normalization options include:
  - None (default). LFQ intensities from MaxQuant or FragPipe have already been normalized with the MaxLFQ [6] algorithm.
  - Quantile normalization. Makes intensity distributions identical in statistical properties.
  - Variance stabilization normalization (VSN). Transforms intensities in such a way that the variance remains almost constant over the whole intensity spectrum.
  - Median centering normalization. Intensities from each sample are scaled in such a way that they all have the same median.

- Differential expression analysis using:
  - limma [7] (default). Moderated t-statistics with empirical Bayes smoothing of protein-wise variance provides increased power.
  - DEqMS [8]. Built on top of limma, includes the number of PSMs or peptides in its variance estimation.

If a pilot experiment without replicates is uploaded fold changes are calculated by subtracting the log-transformed values from the samples in the contrast matrix.

**Missing value imputation**

- min (minimum intensity value): All missing values are replaced by this constant which is useful for pilots.
- normal: imputes each sample from a normal distribution downshifted 1.8 standard deviations from the mean with a width of 0.3 standard deviations.
- global: imputes like the normal option but for the complete data matrix.
- Downshift: (default 1.8).
- Width: (default 0.3).

### Chapter 3

### Input tab

#### 3.1 Upload and Analysis

After successfully uploading and analyzing a dataset, users can download the results in amica's format (Fig. 4a). Additionally, summary information about the uploaded files and a parameter summary of the analysis is outputted. The uploaded experimental design file is depicted as a table (Fig. 4b).

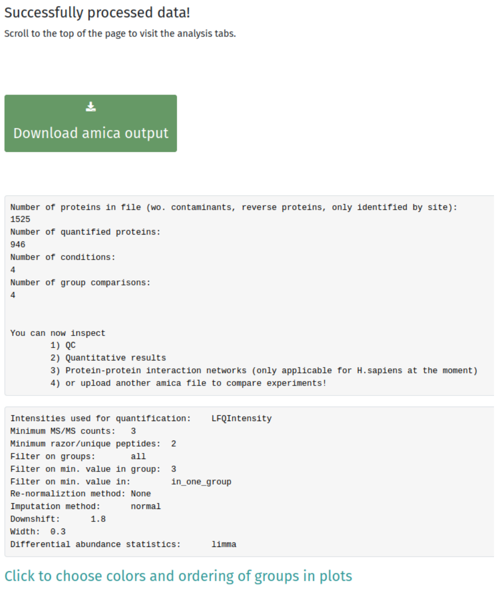

(a) Experimental design table.

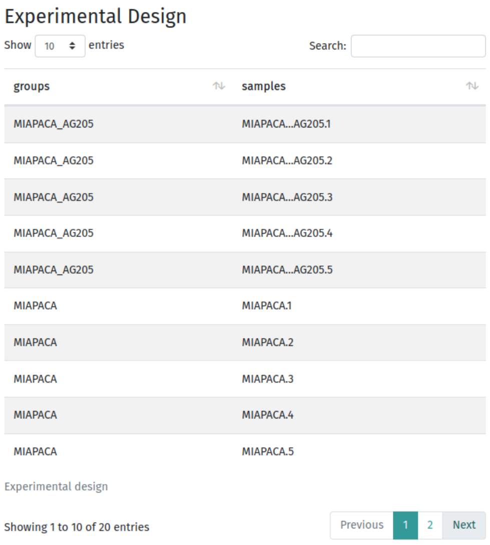

(b) Analysis summary.

Figure 4: Input tab.

#### 3.2 Colors

Color palettes from ColorBrewer [9] can be selected (Fig. 5), after users have successfully uploaded their data.

**Qualitative (groups) colors** are applied to the following visualizations:

- PCA plots
- Box - and barplots (intensities, identified proteins, missing values, CVs)
- Correlation plots group annotations
- Heatmaps group annotations

In addition to the color palettes from ColorBrewer, users can select specific colors for each group by using a color input tool.

**Qualitative (scatter) colors** are applied to the following visualizations:

- Scatter plots
- Volcano - and MA - plots
- Fold-change plots
- PPI networks

**Diverging colors** are applied to the following visualizations:

- Heatmaps (in QC-tab and in Differential abundance tab)
- Correlation plots (in QC-tab and in Compare amica datasets tab)

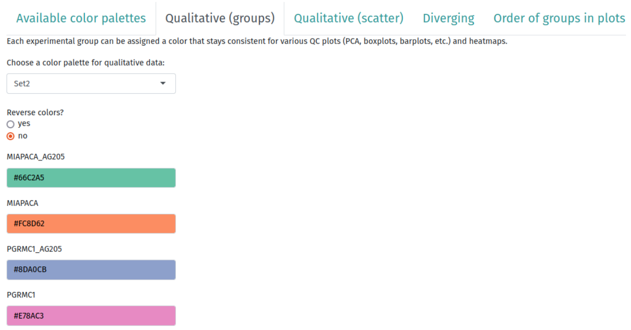

Figure 5: Colors and factors. Clicking on any of the color bars will enable a color picker tool.

#### 3.3 Defining the ordering of groups in visualizations

Users can select the ordering of groups on the x-axis and in legends of plots for QC-plots such as PCA -, bar - and boxplots as well as for profile plots. This feature can also be used to exclude groups from visualizations in the QC tab. If no ordering is provided, these visualizations will be ordered alphabetically.

### Chapter 4

### QC tab

#### 4.1 Plots for different intensities

Various QC output graphics can be generated for different intensity types stored in amica. Users can select from the following intensity types:

- **LFQIntensity** or **Intensity** are intensities that still contain missing values and potential contaminants. These intensities are used to calculate the fraction of missing data or the number of identified protein groups in a sample. If the data was quantified with MaxQuant or FragPipe these intensities are usually already normalized.

- **ImputedIntensity** are normalized and imputed intensities that are also used to calculate differential abundance. If the re-normalization option was selected in the input tab the LFQIntensities were normalized after removing potential contaminants, reverse hits, proteins only identified by site and protein groups that had too few valid values per group.

- **iBAQ** [10] (intensity-based absolute quantification) values are obtained by dividing protein intensities by the number of theoretically observable peptides. This measure correlates well with protein abundance and is used, for example, to calculate the percentage of contamination in a sample (if available).

- **RawIntensity** are non-normalized, summed peptide intensities per protein group.

Principle Component Analysis (PCA, Fig. 6a) is a dimensionality reduction technique that enables the visual inspection of potential clustering in the data. The Pearson correlation coefficient of all samples is visualized as a heatmap (Fig. 6b) and it is a measure of linear correlation. For both plots, users can select which groups should be visualized (if no selection is made, all groups are plotted). An additional color annotation mapping samples to distinct biological groups is available for the correlation plot.

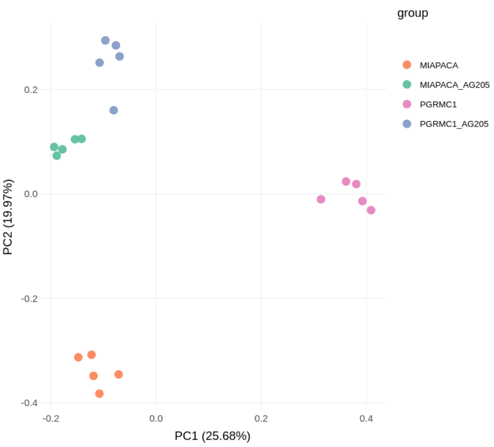

(a) PCA plot.

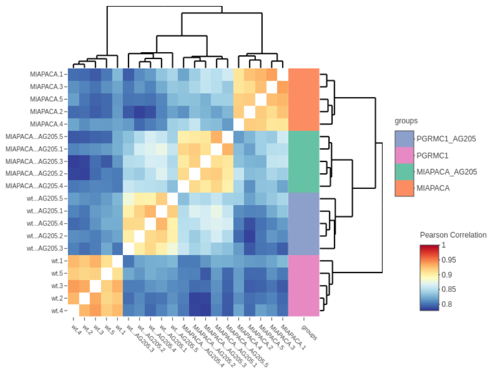

(b) Correlation plot.

Figure 6: Exploratory data analysis.

The distribution of sample intensities can be visualized as box plots (Fig. 7a), or as density plots (Fig. 7b).

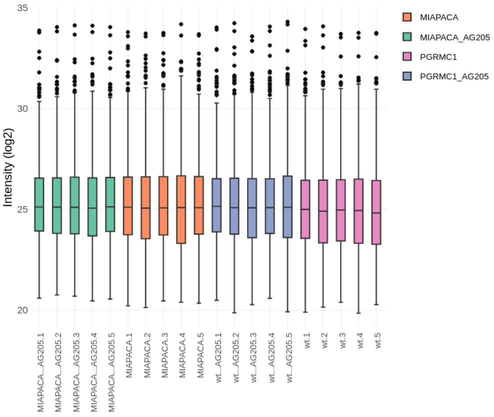

(a) Box plot.

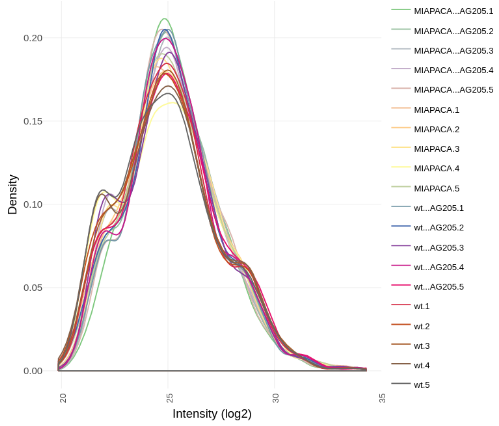

(b) Density plot.

Figure 7: QC plots.

The Coefficient of Variation (CV), shown in in Figure 8a, is the the standard deviation of replicates divided by their mean per protein, which gives an estimate on the reproducibility of the experiment. This plot is only rendered when replicates are available. For scatter plots (Fig. 8b), users can select the intensity type and the sample name to plot, which allows for the visualization of different intensities for the same sample (e.g raw intensity vs. normalized intensity). For the scatter plot, users have the option of plotting a line (no line, straight line or linear regression).

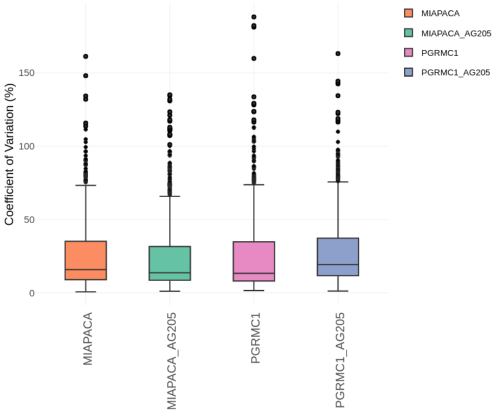

(a) CV plot.

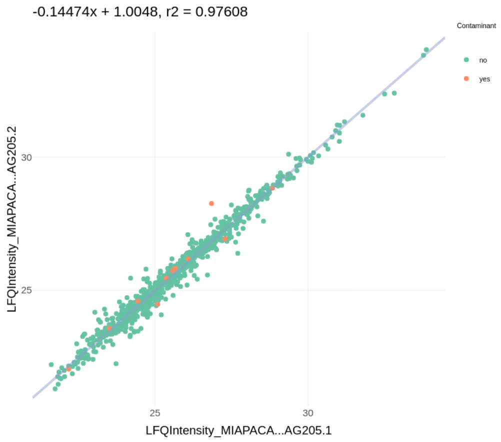

(b) Scatter plot.

Figure 8: QC plots.

#### 4.2 Plots for iBAQ and LFQ intensities

Bar plots of the number of identified proteins (Fig. 9a) and bar plots depicting the percentage of missing values (Fig. 9b) are only shown when `LFQIntensity` - or `Intensity` values are present in the uploaded data.

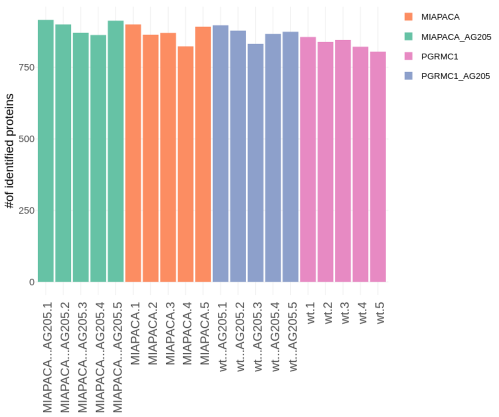

(a) Bar plot of identified proteins.

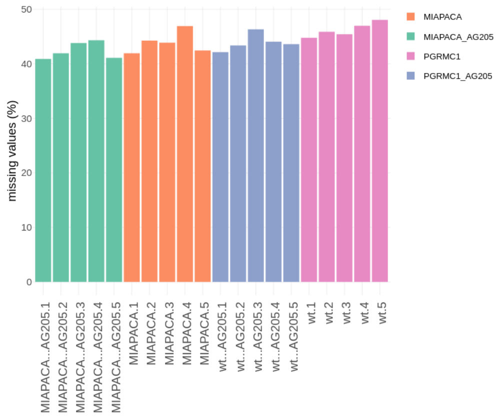

(b) Bar plot of missing values.

Figure 9: Identified proteins and missing value barplots.

A heatmap (Fig. 10a) and a corresponding data table (Fig. 10b) can be inspected to compare the overlap of identified protein groups between samples. Users can select the Jaccard index, the overlap coeffcient or the number of shared protein groups to be displayed in the heatmap. An additional color annotation mapping samples to distinct biological groups is available for the heatmap.

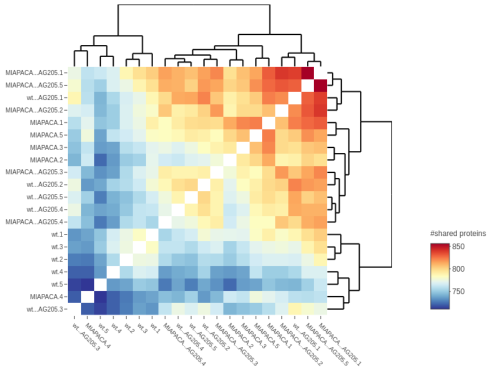

(a) Sample overlap heatmap.

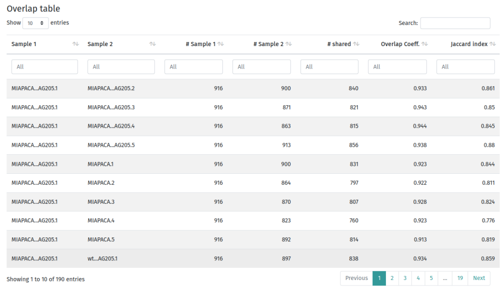

Overlap table

Show 10 entries Search:

| Sample 1 | Sample 2 | # Sample 1 | # Sample 2 | # shared | Overlap Coeff. | Jaccard index |
|---|---|---|---|---|---|---|
| All | All | All | All | All | All | All |
| MIAPACA...AG205.1 | MIAPACA...AG205.2 | 916 | 900 | 840 | 0.933 | 0.861 |
| MIAPACA...AG205.1 | MIAPACA...AG205.3 | 916 | 871 | 821 | 0.943 | 0.85 |
| MIAPACA...AG205.1 | MIAPACA...AG205.4 | 916 | 863 | 815 | 0.944 | 0.845 |
| MIAPACA...AG205.1 | MIAPACA...AG205.5 | 916 | 913 | 856 | 0.938 | 0.88 |
| MIAPACA...AG205.1 | MIAPACA.1 | 916 | 900 | 831 | 0.923 | 0.844 |
| MIAPACA...AG205.1 | MIAPACA.2 | 916 | 864 | 797 | 0.922 | 0.811 |
| MIAPACA...AG205.1 | MIAPACA.3 | 916 | 870 | 807 | 0.928 | 0.824 |
| MIAPACA...AG205.1 | MIAPACA.4 | 916 | 823 | 760 | 0.923 | 0.776 |
| MIAPACA...AG205.1 | MIAPACA.5 | 916 | 892 | 814 | 0.913 | 0.819 |
| MIAPACA...AG205.1 | wt...AG205.1 | 916 | 897 | 838 | 0.934 | 0.859 |

Showing 1 to 10 of 190 entries Previous 1 2 3 4 5 ... 19 Next

(b) Sample overlap table.

Figure 10: Summary of sample overlap.

The percentage of contaminants per sample can be visualized as a bar plot (Fig. 11a), it is calculated using the iBAQ values.

Users can select a sample to inspect the 15 most abundant proteins, the y-axis shows the percentage of iBAQ values for these proteins in that sample (Fig. 11b).

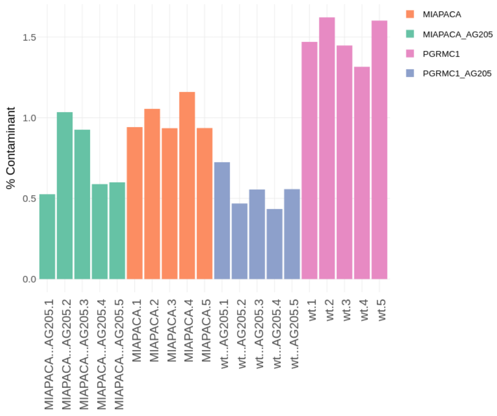

(a) Contamination bar plot.

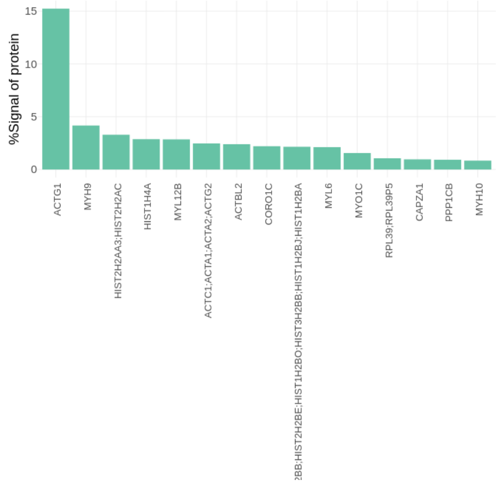

(b) Most abundant proteins bar plot.

Figure 11: Contaminants and most abundant proteins plot.

### Chapter 5

### Differential abundance tab

#### 5.1 Global parameters

The following parameters can be used to subset the data to show only protein groups of interest. The selected parameters are applied to all user-selected group comparisons and include:

- Fold change threshold: While the choice is arbitrary, lower thresholds might result in more false positives. Popular choices are between 1 and 2.
- Significance cutoff:
  - adjusted p-value (recommended)
  - p-value
  - none
- (adj.) p-value threshold: more stringent p-value thresholds can be set, default value is 0.05.
- How to apply fold change threshold
  - absolute: selects significant proteins above and below that negative threshold (e.g if fold change threshold equals 2 'absolute' selects proteins in the ranges [-inf - (-2)] and [2 - inf]).
  - enriched: selects only significant proteins having a fold change greater than the threshold.
  - reduced: selects only significant proteins below the negative fold change threshold.

#### 5.2 Analyzing single group comparisons

In volcano plots (Fig. 12a) the fold change is shown on the x-axis and the -$\log_{10}$ p-value are plotted on the y-axis. Proteins with the biggest quantitative differences are located in the top left and top right corners of the plot. MA plots (Fig. 12b) show the fold change on the x-axis and the average intensities from the selected groups for each protein on the y-axis. A lasso select option, which becomes visable when users move their mouse over the plots, is available for both volcano - and MA - plots. The lasso select options highlights selected proteins in the plot by showing their gene names next to their data point. User-defined colors for non-significant and significant proteins can be selected for both plots.

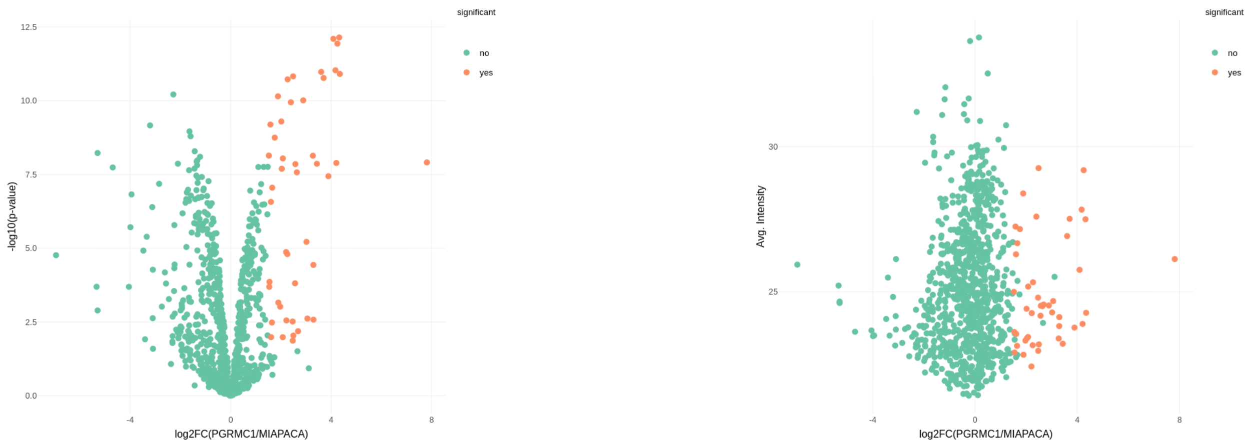

(a) Volcano plot.

(b) MA plot.

Figure 12: Differential abundance plots.

#### 5.3 Analyzing multiple group comparisons

Users can visualize the overlap of significant proteins from multiple selected group comparisons in Euler diagrams (Fig. 13a) and UpSet plots [11] (Fig. 13b). The dots in the UpSet plot show which sets are being compared. A dot not connected to another dot shows the number of proteins specific to that group. The top bar plot depicts the number of intersecting proteins, and the bar plot on the side shows how many proteins are differentially abundant in the comparison.

For both plots, users can change the labels of the selected group comparisons. Euler plots are only available for up to five group comparisons. User-defined colors can be chosen in the Euler diagram.

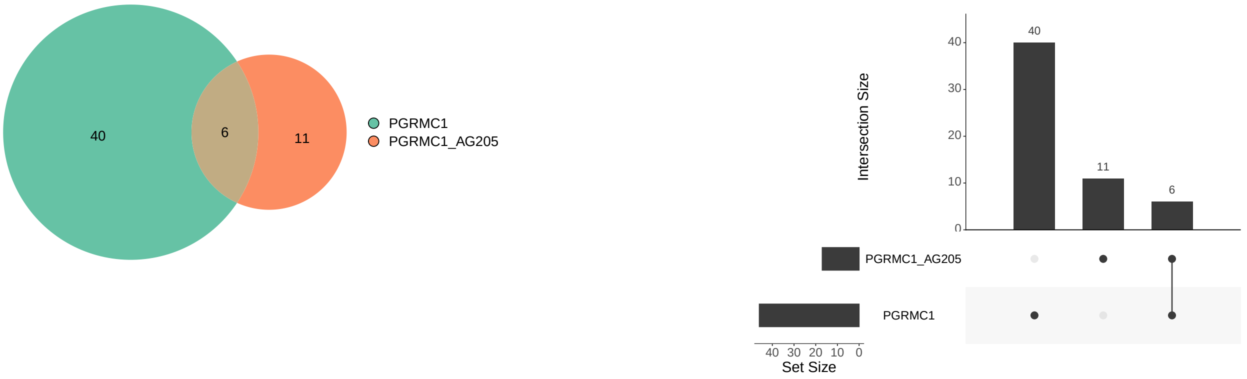

(a) Euler plot.

(b) Upset plot.

Figure 13: Visualization of multiple selected group comparisons.

#### 5.4 Data table

Significant proteins are displayed together with their differential expression statistics in a data table that can be further filtered. Users can for example use regular expressions (e.g type in a query like "`ProteinA|ProteinB|ProteinC`" in the Gene.names column) to select only proteins of interest or apply additional column filters (Fig. 14). When multiple group comparisons are selected, users can subset the proteins that are specific to one group comparison only (e.g by writing 'yes' in one significant column and 'no' in the other column.)

Subsequent visualizations (heatmap, fold change plot, PPI network) and Over-Representation Analysis (ORA) will be computed on the proteins in the output table.

| Gene.names | significant_PGRMC1__vs__MIAPACA | logFC_PGRMC1__vs__MIAPACA | P.Value_PGRMC1__vs__MIAPACA |
|---|---|---|---|
| 1\|SPTBN1\|SPTAN1\|FSCN1\|SSH1\|MYO18A\|CNN2\|CORO1B\|ANLN | All | All | All |
| ARPC1B | yes | 2.4702 | 0.01353 |
| ACTN4 | yes | 4.2487 | 1.160e-12 |
| FLNB | yes | 2.3981 | 1.130e-10 |
| MYO1D | yes | 2.5744 | 1.403e-8 |
| ACTN1 | yes | 4.3205 | 7.175e-13 |
| FLNA | yes | 4.0877 | 7.987e-13 |
| IQGAP1 | yes | 1.8830 | 7.089e-11 |
| SPTBN1 | yes | 3.6026 | 1.055e-11 |
| SPTAN1 | yes | 3.7002 | 1.691e-11 |
| FSCN1 | yes | 2.2058 | 0.00001342 |

Showing 1 to 10 of 15 entries (filtered from 46 total entries)

There are 46 proteins in your selection. After filtering the output table 15 proteins remain for subsequent visualizations. Remove the filters in the table to visualize all proteins.

Figure 14: A data table subsetted using regular expressions. Below the data table a text message informs the user how many proteins remain after subsetting the data.

#### 5.5 Plots dependent on the data table

##### Heatmap

Users can select different options for the color gradient to plot, such as values from imputed intensities or z-score transformed intensities from rows or columns. Dendrograms, as well as row and column annotations can be disabled. Additionally, a color code for groups can be added to the column dendrogram. An example heatmap is depicted in Figure 15.

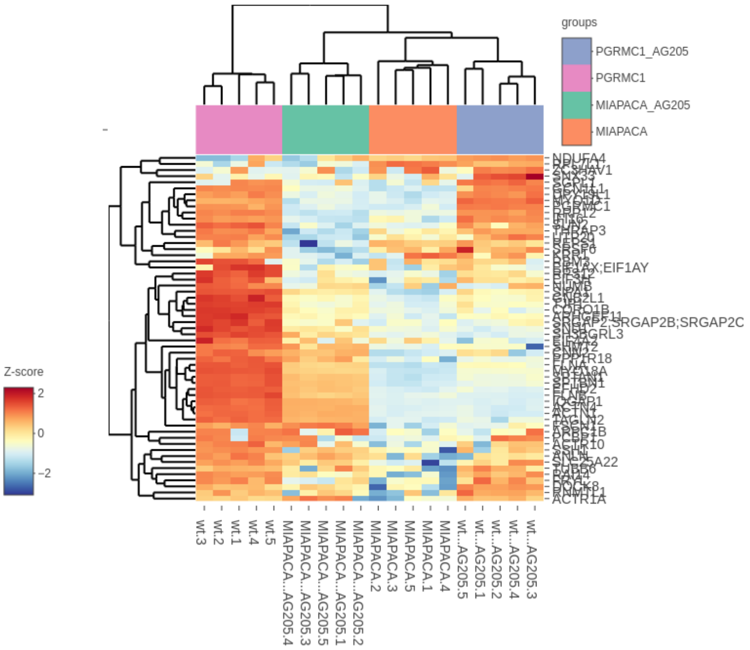

Figure 15: Interactive heatmap.

##### Dot plot

Dot plots (Fig. 16) integrate quantitive information together with log2 fold changes and (adj.) p-values into one visualization. Proteins are displayed as dots, with their circle size corresponding to relative abundance (average intensities or log2 fold changes can be selected). Log2 fold changes are mapped as color gradients on the dots. When proteins are clustered based on fold changes, users have the option to display proteins with a positive fold change only. Dot sizes and a minimum and maximum value for the log2FC color range can be selected. Users can cluster the rows and columns with multiple distance metrics (canberra, euclicadean, maximum, manhattan, binary and minkowski), as well as with a variety of clustering methods (complete, average, single, Ward, median and centroid).

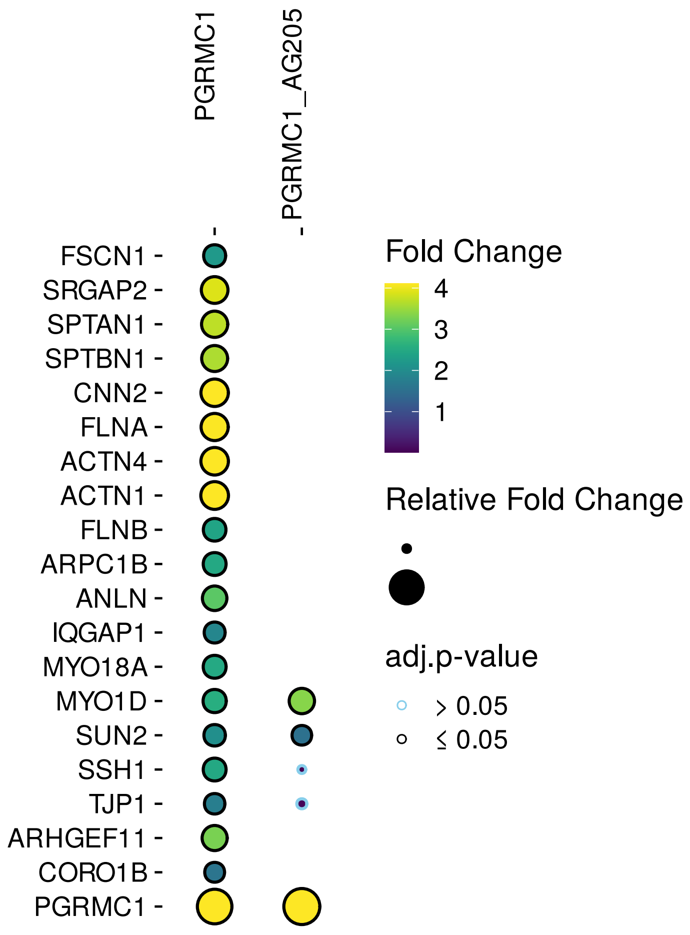

Figure 16: Interactive heatmap.

#### Protein-Protein Interaction (PPI) Network

PPI Networks from the IntAct database [12] (downloaded on October 2021) are retrieved for the proteins in the data table. All interactions are derived from literature curation or direct user submissions and only contain experimental evidence from low - and high - throughput experiments. The edge widths between PPIs correspond to the MI score [13], which evaluates the confidence in the interaction; the higher the edge width, the higher the interaction confidence.

Subcellular location predictions are retrieved from the humancellmap database [14] (downloaded on October 2021) and can be highlighted in the network. The humancellmap database provides two different subcellular localization predictions, one resulting from Spatial analysis of functional enrichment (SAFE) and one resulting from non-negative matrix factorization (NMF). A summarized subcellular localization from NMF predictions can be mapped onto the nodes (Fig. 18b), predictions from SAFE and NMF can be seen in a node data table.

The network can be downloaded as html file or in gml format (including subcellular localization information).

**PPI network for single group comparisons** For single group comparisons, a fold change color gradient is mapped onto the nodes in the network (Fig. 17).

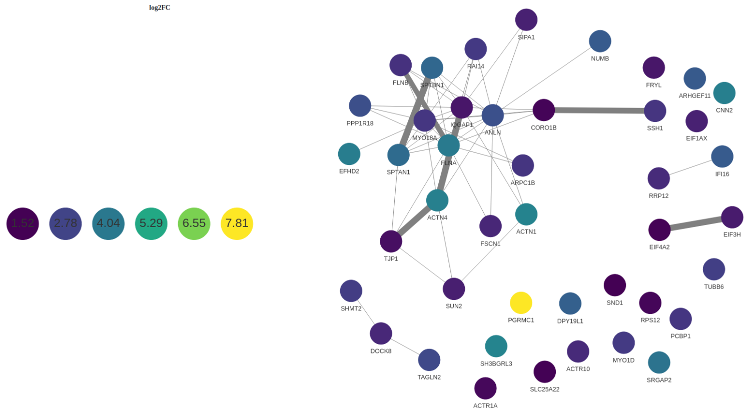

Figure 17: interactive PPI network.

**PPI network for multiple group comparisons** For multiple group comparisons the network contains two types of edges: (1) edges corresponding to PPI from the IntAct database and (2) edges between group comparisons and their significant proteins (Fig. 18a).

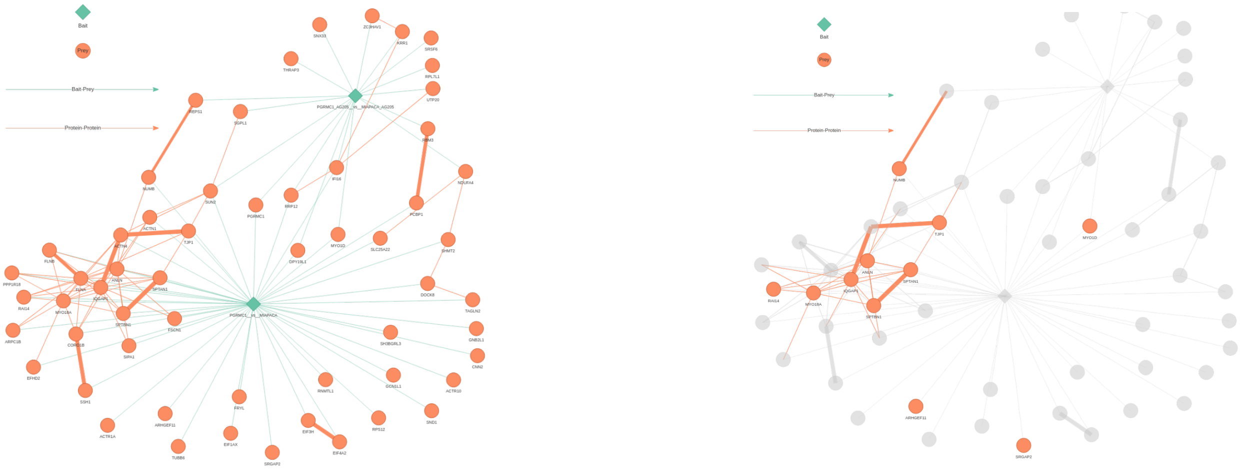

(a) PPI network for multiple group comparisons.

(b) Human Cell Map predictions (plasma membrane, cell junction) mapped on the nodes.

Figure 18: PPI network for multiple selected group comparisons.

#### Fold-change plot

Fold-change plots (Fig. 19) are useful to compare proteins across different comparisons, data points are colored based on their significance to the comparison (e.g whether they are significant in both comparisons, in one comparison only, or in none). For the fold-change plot, users have the option of plotting a line (no line, straight line or linear regression). A lasso select options is available to highlight selected data points and users can change the name of the group comparison that is shown in the legend.

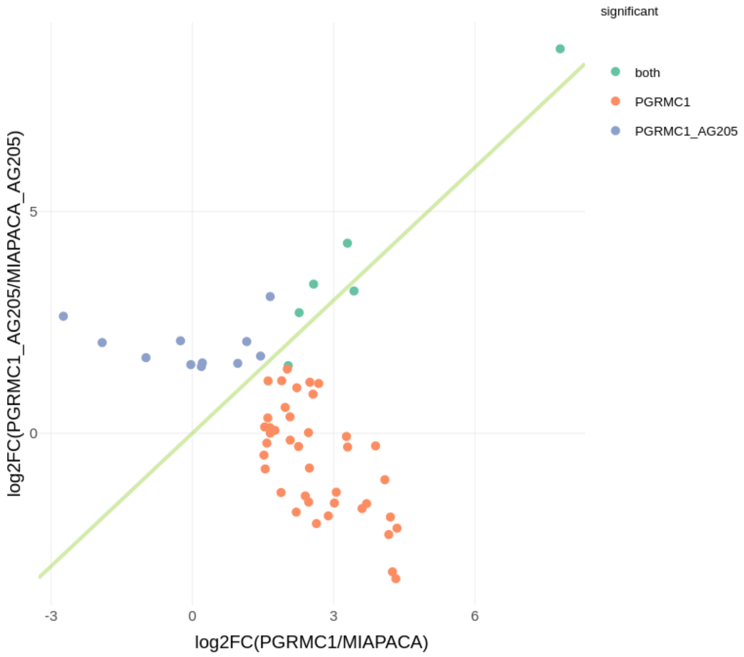

Figure 19: Fold change plot.

#### 5.6 Profile plots

For profile plots, users can select a protein to plot from all quantified proteins. Users can select the mean of imputed intensities together with the standard error of the mean (if replicates are available), violin plots or only the individual data points to be plotted (Fig. 20a) for all groups. Summary statistics from the differential expression analysis are shown below the profile plot (Fig. 20b), allowing for the immediate inspection of proteins of interest.

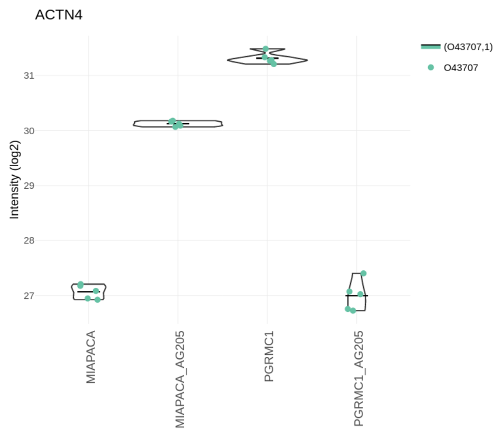

(a) Profile plot.

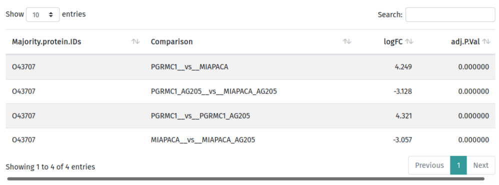
Show 10 entries Search:

| Majority.protein.IDs | Comparison | logFC | adj.P.Val |
|---|---|---|---|
| O43707 | PGRMC1__vs__MIAPACA | 4.249 | 0.000000 |
| O43707 | PGRMC1_AG205__vs__MIAPACA_AG205 | -3.128 | 0.000000 |
| O43707 | PGRMC1__vs__PGRMC1_AG205 | 4.321 | 0.000000 |
| O43707 | MIAPACA__vs__MIAPACA_AG205 | -3.057 | 0.000000 |

Showing 1 to 4 of 4 entries Previous 1 Next

(b) Summary statistics.

Figure 20: PPI network for multiple selected group comparisons.

#### 5.7 Over-Representation Analysis (ORA)

ORA is a method to determine whether pre-defined gene sets (e.g from pathway - or ontology databases) are over-represented in a subset of the significantly differentially abundant proteins. This approach assists in unveiling changes in the underlying biology. Users can set options to show the gene names corresponding to a functional term or to also include non-significant terms in the output table (Fig. 21). Furthermore, users can select all organisms present in gprofiler [15] (organism list downloaded on October 2021) and can specify multiple databases to include in the analysis:

- Gene Ontology (GO)
  - GO:MF (Molecular Function)
  - GO:CC (Cellular Component)
  - GO:BP (Biological Process)
- REAC (Reactome): pathway database
- KEGG: pathway database
- CORUM: mammalian protein complexes database
- WP (Wiki Pathways): pathway database
- TF (TRANSFAC): database of eukaryotic transcription factors
- MIRNA (miRTarBase): database for miRNA targets
- HPA (Human protein Atlas): database for tissue specificity
- HP (Human Phenotype Ontology): database for human disease phenotypes

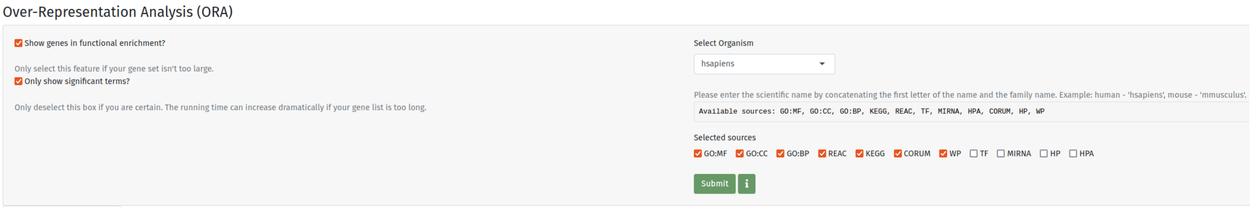

Figure 21: ORA parameters.

An overview of all selected sources is shown in a Manhattan plot (Fig. 22a) which shows the -$\log_{10}$ p-value for every term as a circle, whose size corresponds to the number of genes in that term.

For each source database, a bar plot (Fig. 22b) depicting the -$\log_{10}$ p-value of the functional enrichment can be generated. Users can change the color of the bars and specify the maximum number of terms to be included in the plot.

The results of the ORA are also outputted in a downloadable data table (Fig. 23).

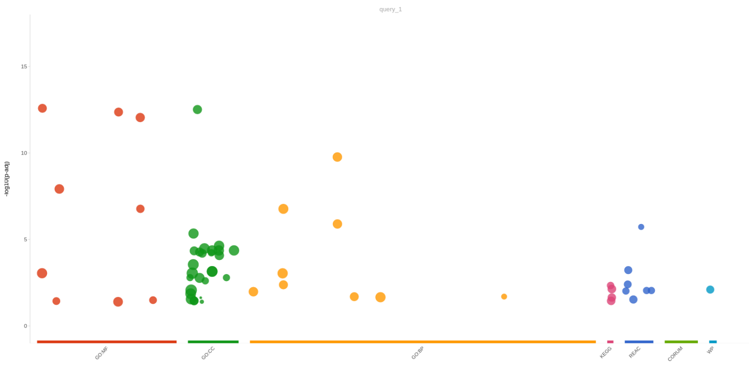

(a) Manhattan plot.

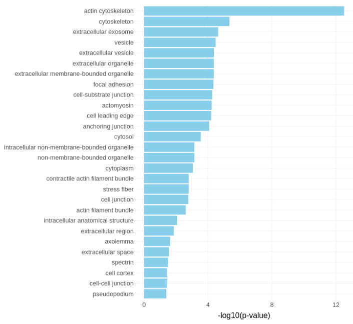

(b) ORA barplot.

Figure 22: ORA results.

Download results

Show 10 entries

Search:

| source | term_id | term_name | p_value | term_size | intersection_size | intersection |
|---|---|---|---|---|---|---|
| All | All | All | All | All | All | All |
| GO:MF | GO:0003779 | actin binding | 2.62e-13 | 320 | 15 | ARPC1B,ACTN4,FLNB,MYO1D,ACTN1,FLNA,IQGAP1,SPTBN1,SPTAN1,FSCN1,SSH1,MYO18A,CNN2,CORO1B,ANLN |
| GO:CC | GO:0015629 | actin cytoskeleton | 3.10e-13 | 443 | 16 | ARPC1B,ACTN4,FLNB,MYO1D,ACTN1,FLNA,IQGAP1,ACTR1A,SPTBN1,SPTAN1,FSCN1,MYO18A,CNN2,CORO1B,ANLN,ACTR10 |
| GO:MF | GO:0045296 | cadherin binding | 4.33e-13 | 331 | 15 | FLNB,FLNA,TAGLN2,IQGAP1,NUMB,SPTBN1,TJP1,SPTAN1,PCBP1,FSCN1,SND1,EFHD2,CNN2,CORO1B,ANLN |
| GO:MF | GO:0050839 | cell adhesion molecule binding | 8.95e-13 | 524 | 17 | ACTN4,FLNB,ACTN1,FLNA,TAGLN2,IQGAP1,NUMB,SPTBN1,TJP1,SPTAN1,PCBP1,FSCN1,SND1,EFHD2,CNN2,CORO1B,ANLN |
| GO:BP | GO:0030029 | actin filament-based process | 1.72e-10 | 718 | 17 | ARHGEF11,ARPC1B,ACTN4,SRGAP2,FLNB,MYO1D,ACTN1,FLNA,IQGAP1,TJP1,FSCN1,SSH1,MYO18A,CNN2,CORO1B,ANLN,SUN2 |
| GO:MF | GO:0008092 | cytoskeletal protein binding | 1.21e-8 | 802 | 16 | ARPC1B,ACTN4,FLNB,MYO1D,ACTN1,FLNA,IQGAP1,SPTBN1,SPTAN1,FSCN1,SSH1,MYO18A,CNN2,CORO1B,ANLN,SUN2 |
| GO:MF | GO:0051015 | actin filament binding | 1.70e-7 | 178 | 9 | ARPC1B,ACTN4,MYO1D,ACTN1,FLNA,IQGAP1,FSCN1,MYO18A,CORO1B |

Figure 23: ORA table.

### Chapter 6

### Compare amica datasets tab

#### 6.1 Upload

amica allows users to upload a second amica file to be compared with the current data input. Parameters to merge the uploaded input file with the currently loaded in data input are shown in Figure 24. Users can specify the key column which should be used to merge the data (either the unique protein id or the gene name). A suffix for the original input and the uploaded input can also be specified that adds this suffix to all column names of the corresponding inputs. Lastly, a pattern can be specified that splits every id in the key column.

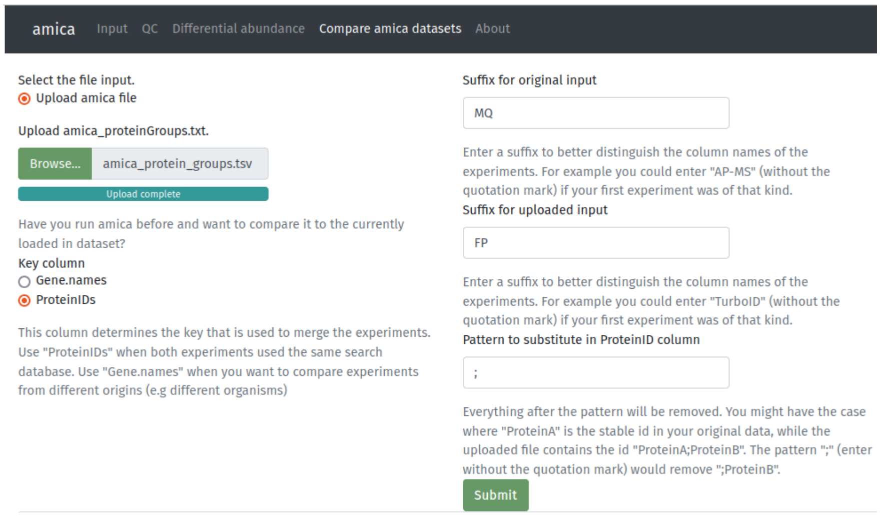

Figure 24: Second amica file upload section.

#### 6.2 Correlation analysis

After succesfully uploading a second amica file users can download the merged datasets. Different intensities can be selected to be correlated in scatter plots (Fig. 25a), where users can also select two samples to compare as well as different options for fitting a line through the data (no line, straight line or linear regression). Output graphics also include correlation plots (Fig. 25b) between all samples from both experiments.

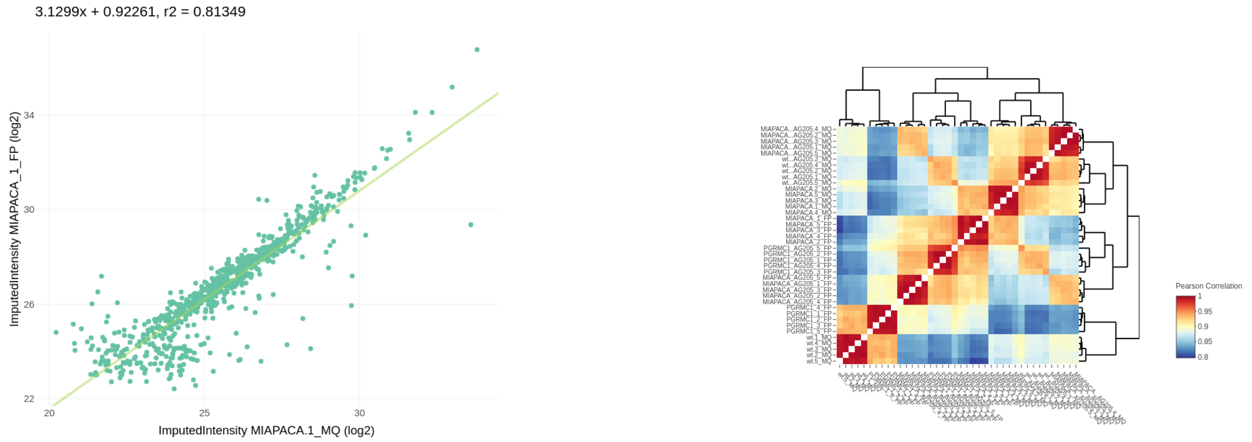

(a) Scatter plot. (b) Correlation plot of LFQ intensities.

Figure 25: Correlation analysis of the combined dataset.

#### 6.3 Analyze combined dataset in Differential abundance tab

At the top of the differential abundance tab a select box becomes visable that lets users choose between the original data (`original_data`) and the combined dataset (`multiple_amica_data`). After selecting `multiple_amica_data` and pressing the Submit button, the complete functionality of the differential abundance tab (with the exception of heatmaps and profile plots), becomes available for the combined dataset and the differential abundance tab can be be used as for a regular upload.

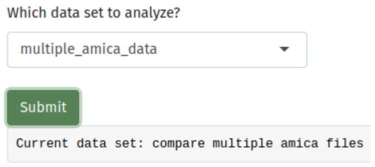

Figure 26: Data selection box at the top of the differential abundance tab.

### Chapter 7

### Tutorials

#### 7.1 How to convert analyzed data into amica format

If you want to upload data into amica that has already been analyzed in a different tool or context (e.g data from RNA-Seq) you need to change the column names of your file into amica's column names.

The following toy dataset has a unique id, a gene name, some expression values and statistics from a differential expression analysis:

| uniqueID | Gene | logExpr_sample_1 | logExpr_sample_2 | ... | logExpr_sample_n | pval_trtmt/ctrl | padj_trtmt/ctrl | logfc_trtmt/ctrl |
|---|---|---|---|---|---|---|---|---|
| id_1 | Gene_1 | 30 | 30.5 | ... | 28.2 | 0.00012 | 0.002 | 1.7 |
| id_2 | Gene_2 | 28.6 | 28.5 | ... | 26.9 | 0.0002 | 0.003 | 1.68 |
| ... | ... | ... | ... | ... | ... | ... | ... | ... |
| id_p | Gene_p | 20 | 20.3 | ... | 18 | 0.99 | 0.99 | -0.02 |

The `uniqueID` column needs to be renamed into `Majority.protein.IDs`,

the `Gene` column into `Gene.names`,

and all `logExpr_` prefixes need to be replaced by `ImputedIntensity_`
(e.g `ImputedIntensity_sample_1`, `ImputedIntensity_sample_2`, ..., `ImputedIntensity_sample_n`)

Columns containing the results from the differential expression analysis (`pval_trtmt/ctrl`, `padj_trtmt/ctrl`, `logfc_trtmt/ctrl`) need to be adapted in such a way that they contain the correct prefixes and the `__vs__` - infix.

`pval_trtmt/ctrl` has to be changed to `P.Value_trtmt__vs__ctrl`,
`padj_trtmt/ctrl` to `adj.P.Val_trtmt__vs__ctrl` and
`logfc_trtmt/ctrl` to `logFC_trtmt__vs__ctrl`.

Furthermore, you could specify a `quantified` column that contains for each entry a "+" if it has been quantified, else it needs to be left empty. If no `quantified` column is provided, amica automatically considers all entries quantified that do not contain missing values in the ImputedIntensity and `__vs__` - infix columns.

The columns should now have following names:

| |
|---|
| Majority.protein.IDs |
| Gene.names |
| ImputedIntensity_sample_1 |
| ImputedIntensity_sample_1 |
| ... |
| ImputedIntensity_sample_n |
| P.Value_trtmt__vs__ctrl |
| adj.P.Val_trtmt__vs__ctrl |
| logFC_trtmt__vs__ctrl |

Save this file as a tab-separated txt file (you can choose a file name of your choice, the output name of amica is by default amica_protein_groups.txt).

Finally, we need to create a tab-separated experimental design that assigns the samples to their appropriate group. Here, it is important to link the groups to the p-value and fold change columns of the group comparison infixes (e.g `logFC_trtmt__vs__ctrl` corresponds to the group comparison `trtmt` vs `ctrl`). All groups from the group comparison infixes need to be defined in the experimental design. If you have multiple 'Intensity' - prefixes in your amica file, it is important that all of them have the same number of samples. The sample names in the samples column of the design need to match the column names of the input file in the order of the input file.

| groups | samples |
|---|---|
| trtmt | sample_1 |
| trtmt | sample_2 |
| trtmt | sample_3 |
| ctrl | sample_4 |
| ctrl | sample_5 |
| ctrl | sample_6 |

Save this file as a tab-separated txt file (you can choose a file name of your choice). Now you can upload both files and explore your data in amica.

#### 7.2 How to use the differential abundance tab

These small examples have been produced with the provided example data set [1], an interaction proteomics study focusing on PGRMC1, a protein from the MAPR family with a range of cellular functions. In this study, MIA PaCa-2 cells were stably transfected with a PGRMC1-HA plasmid and Co-IPs of PGRMC1 interacting proteins were isolated from cells expressing PGRMC1-HA, as well as from non-transfected parental MIA PaCa-2 cells as a negative control, with or without AG-205 treatment (a PGRMC1-specific inhibitor) [1]. As the example data contains AP-MS data, we set the selection parameter to "enriched" (Fig. 27) to retrieve differentially abundant proteins compared against the control.

##### 7.2.1 Use case 1: Single group comparison

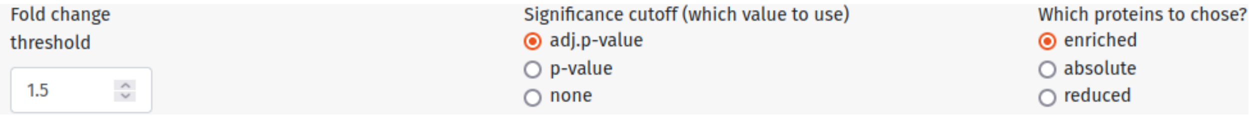

Figure 27: Global parameters for the example data set.

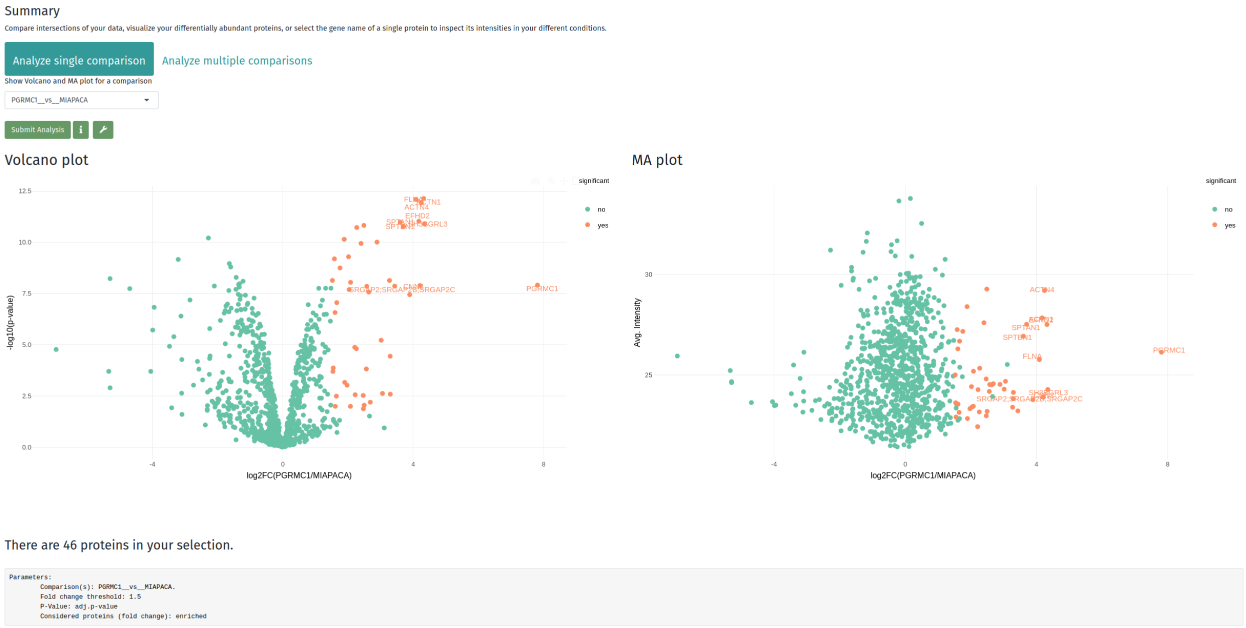

Figure 28: Volcano - and MA - plots.

When you hover over the volcano - or MA - plot you can see features to manipulate the plot. When we utilize the select box or lasso tool we can annotate the highest enriched proteins, as seen in the figure above (Fig. 28).

Further information on most plots can be acquired when you press the "info" icon. Plot parameters can be changed when pressing the "wrench" icon and can be saved upon hovering over the plot and clicking on the "camera" icon.

##### 7.2.2 Advanced queries: Visualize proteins from functional term in ORA

An over-representation analysis was performed utilizing the 46 enriched proteins from the comparison PGRMC1__vs__MIAPACA. The "Show genes in functional enrichment?" button was selected (Fig. 29).

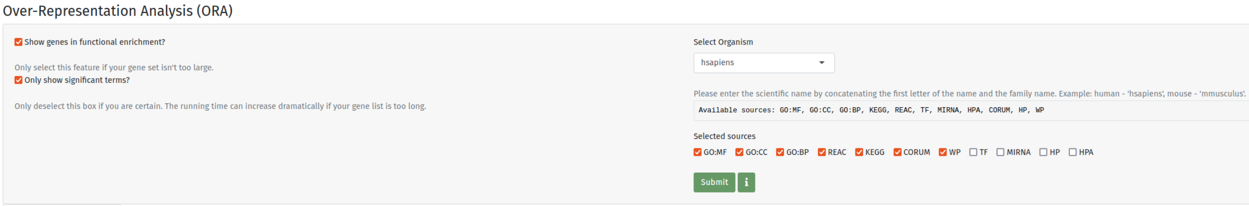

Figure 29: ORA parameters.

Sorting the output table from most significant p-value to least significant we find the term "actin binding" on top of the list (Fig. 30). 15 of the enriched proteins are annotated with this term.

Download results

Show 10 entries

Search:

| source | term_id | term_name | p_value | term_size | intersection_size | intersection |
|---|---|---|---|---|---|---|
| All | All | All | All | All | All | All |
| GO:MF | GO:0003779 | actin binding | 2.62e-13 | 320 | 15 | ARPC1B,ACTN4,FLNB,MYO1D,ACTN1,FLNA,IQGAP1,SPTBN1,SPTAN1,FSCN1,SSH1,MYO18A,CNN2,CORO1B,ANLN |
| GO:CC | GO:0015629 | actin cytoskeleton | 3.10e-13 | 443 | 16 | ARPC1B,ACTN4,FLNB,MYO1D,ACTN1,FLNA,IQGAP1,ACTR1A,SPTBN1,SPTAN1,FSCN1,MYO18A,CNN2,CORO1B,ANLN,ACTR10 |
| GO:MF | GO:0045296 | cadherin binding | 4.33e-13 | 331 | 15 | FLNB,FLNA,TAGLN2,IQGAP1,NUMB,SPTBN1,TJP1,SPTAN1,PCBP1,FSCN1,SND1,EFHD2,CNN2,CORO1B,ANLN |
| GO:MF | GO:0050839 | cell adhesion molecule binding | 8.95e-13 | 524 | 17 | ACTN4,FLNB,ACTN1,FLNA,TAGLN2,IQGAP1,NUMB,SPTBN1,TJP1,SPTAN1,PCBP1,FSCN1,SND1,EFHD2,CNN2,CORO1B,ANLN |
| GO:BP | GO:0030029 | actin filament-based process | 1.72e-10 | 718 | 17 | ARHGEF11,ARPC1B,ACTN4,SRGAP2,FLNB,MYO1D,ACTN1,FLNA,IQGAP1,TJP1,FSCN1,SSH1,MYO18A,CNN2,CORO1B,ANLN,SUN2 |
| GO:MF | GO:0008092 | cytoskeletal protein binding | 1.21e-8 | 802 | 16 | ARPC1B,ACTN4,FLNB,MYO1D,ACTN1,FLNA,IQGAP1,SPTBN1,SPTAN1,FSCN1,SSH1,MYO18A,CNN2,CORO1B,ANLN,SUN2 |
| GO:MF | GO:0051015 | actin filament binding | 1.70e-7 | 178 | 9 | ARPC1B,ACTN4,MYO1D,ACTN1,FLNA,IQGAP1,FSCN1,MYO18A,CORO1B |

Figure 30: ORA table.

All visualizations (heatmap, fold change plot and PPI network) work only on the proteins selected in the above output table. We can filter that table to show proteins only annotated with our term of interest. The output table can parse "regular expressions", so all we need to do is to copy-paste the comma-delimited gene names into a text editor (or text processing tool like MS Word) and replace all commas with vertical line symbols ("|" which is the logical "or" operator) with the "Find and Replace" tool (Fig. 31):

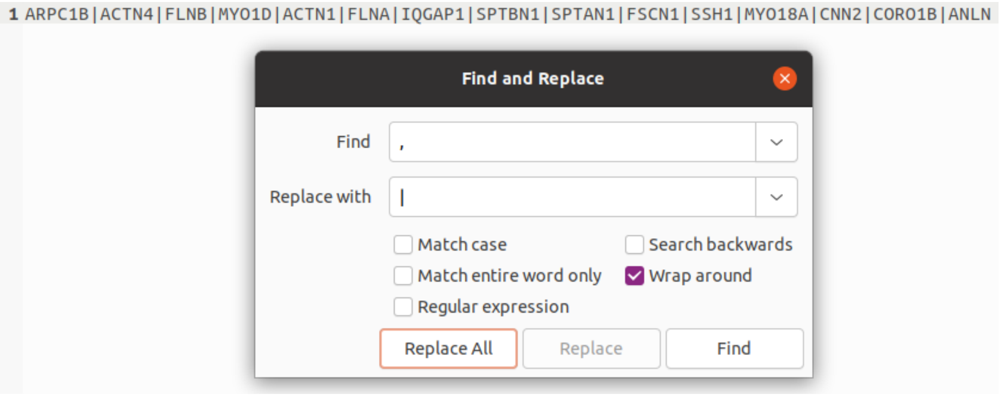

Figure 31: Search and replace.

We can now paste the vertical line-delimited gene names into the "Gene.names" search bar in the output table and successfully subset the data table to show only our proteins of interest (Fig. 32):

| Gene.names | significant_PGRMC1__vs__MIAPACA | logFC_PGRMC1__vs__MIAPACA | P.Value_PGRMC1__vs__MIAPACA |
|---|---|---|---|
| 1\|SPTBN1\|SPTAN1\|FSCN1\|SSH1\|MYO18A\|CNN2\|CORO1B\|ANLN | All | All | All |
| ARPC1B | yes | 2.4702 | 0.01353 |
| ACTN4 | yes | 4.2487 | 1.160e-12 |
| FLNB | yes | 2.3981 | 1.130e-10 |
| MYO1D | yes | 2.5744 | 1.403e-8 |
| ACTN1 | yes | 4.3205 | 7.175e-13 |
| FLNA | yes | 4.0877 | 7.987e-13 |
| IQGAP1 | yes | 1.8830 | 7.089e-11 |
| SPTBN1 | yes | 3.6026 | 1.055e-11 |
| SPTAN1 | yes | 3.7002 | 1.691e-11 |
| FSCN1 | yes | 2.2058 | 0.00001342 |

Showing 1 to 10 of 15 entries (filtered from 46 total entries)

There are 46 proteins in your selection. After filtering the output table 15 proteins remain for subsequent visualizations. Remove the filters in the table to visualize all proteins.

Figure 32: Use Regular Expressions to subset data in data table.

Below the table there is now a text message telling us that the original table has been filtered and that only the remaining proteins are used in subsequent visualizations. As an example you can now observe how the selected proteins compare across different group comparisons in a heatmap or fold change plot.

##### 7.2.3 Use case 2: Multiple group comparisons

The example data contains Co-IPs of PGRMC1 in untreated MIA PaCa-2 cells and MIA PaCa-2 cells treated with AG-205. In this example, we are interested in whether some prey proteins of PGRMC1 are sensitive to AG-205.

When we select the "Analyze multiple comparisons" tab pill we can select the two comparisons of the bait versus the negative controls ( `PGRMC1__vs__MIAPACA`, and `PGRMC1_AG205__vs__MIAPACA_AG205`, Fig. 33):

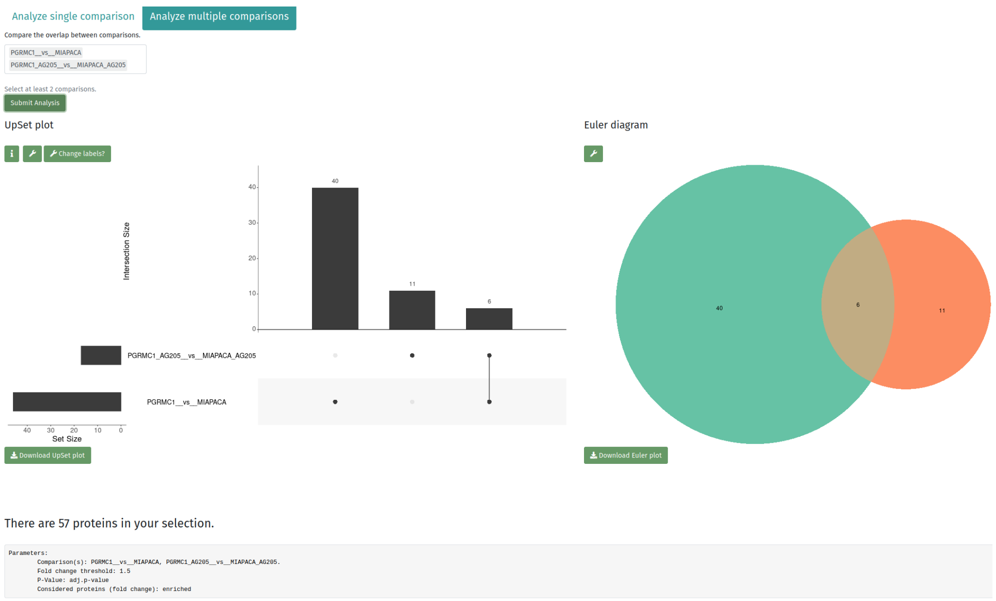

Figure 33: UpSet plot and Euler diagram show that the number of prey proteins of PGRMC1 decreases upon AG-205 treatment.

Scrolling down we can evaluate the quantitative changes of prey proteins with and without AG-205 treatment in a fold-change plot (Fig. 34):

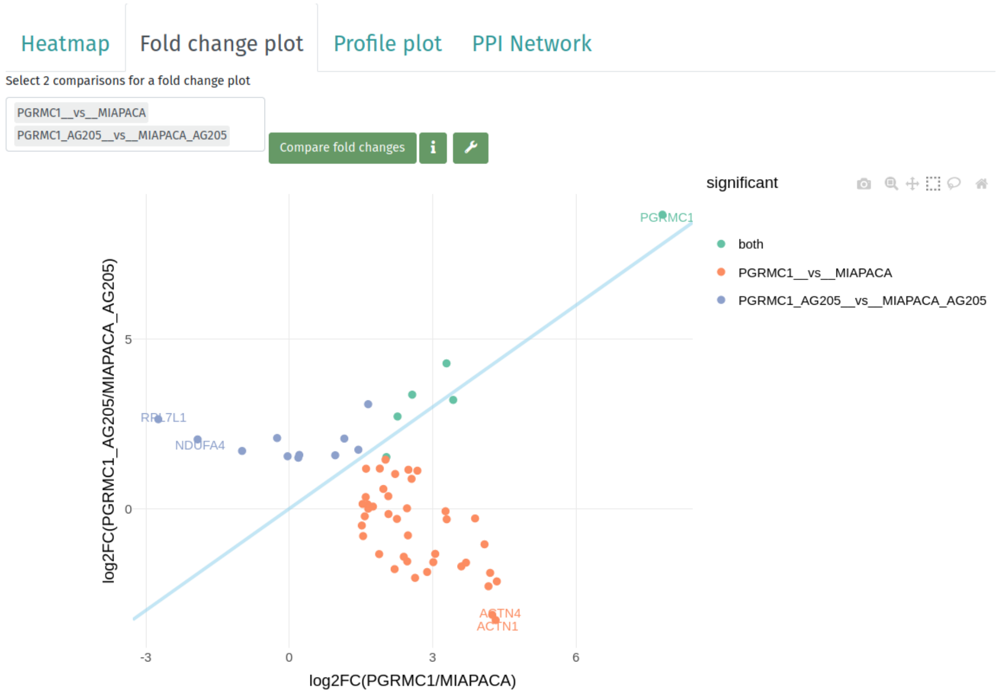

Figure 34: Fold-change plot of PGRMC1 compared against cellular background. Among the highest quantitative changes of AG-205-sensitive proteins are ACTN1 and ACTN4, proteins associated with the actin cytoskeleton.

#### 7.3 How to integrate amica's network output into Cytoscape

Cytoscape [16] is an open source software platform for visualizing complex networks and integrating these with any type of attribute data. The software has many options for analyzing and visualizing networks and is very well documented (you can find very useful tutorials here: `https://github.com/cytoscape/cytoscape-tutorials/wiki`).

We can import amica's PPI networks into Cytoscape for better visualizations. Just download the `amica_specificity_network.gml` file in the Network section in the Differential abundance tab.

Download Cytoscape from `https://cytoscape.org/download.html` if you don't have it already, follow the installation guideline.

Open Cytoscape and import the file `amica_specificity_network.gml`:

```
File > Import > Network from File
```

To produce a layout select any of the layouts in the Layout tab in the main menu:

```
Layout > Apply Preferred Layout
```

We can integrate all the information in the gml file by clicking on the Style tab in the left side bar, where you find the adjustable Node and Edge properties. For each property, styles are defined in columns Def. (default), Map. (mapping), and Byp. (bypass) (Fig. 35).

The first thing we need to do is to show a label for each node. This can be done by clicking on the Map. button of the "Label" property. Select for the Column "label" and for Mapping Type "Passthrough Mapping".

##### 7.3.1 Networks from single group comparisons

The same method can be applied to create a mapping for the logFC column for which we can create a "Continuous Mapping" to create a color bar. Node Size and Shape can also be changed, for that you just have to click on the left "Default" button of these properties. If you want to create circular shapes you have to tick the "lock node width and height" button at the bottom of the Node Style minipage.

When applying all these mappings your Node Style tab should look similar to this one:

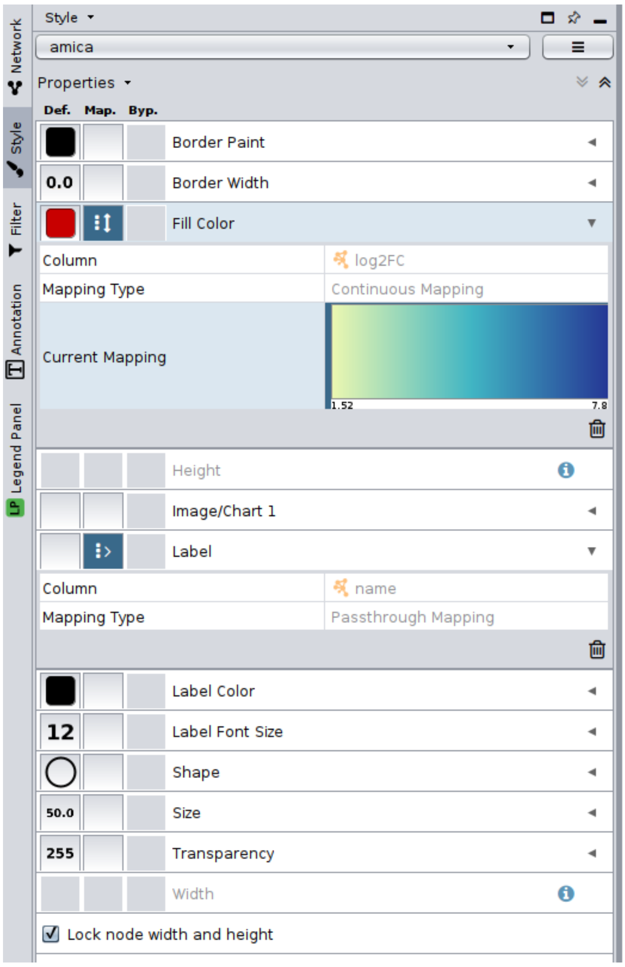

Figure 35: Node style.

To create edge styles click on the Edge tab at the bottom of the Style page. Similarly to the Node Styles, you can change edge properties. One potentially useful feature would be to map the PPI MI-Score to the edge width with a continuous or discrete mapping.

After applying the column mappings for the group comparison `PGRMC1__vs__MIAPACA` from the example data (enriched proteins, log2FC $\geq$ 1.5 and adj.p-value $\leq$ 0.05), we end up with the network like the one shown in Figure 36 (the edge width legend was added manually in inkscape, `https://inkscape.org`):

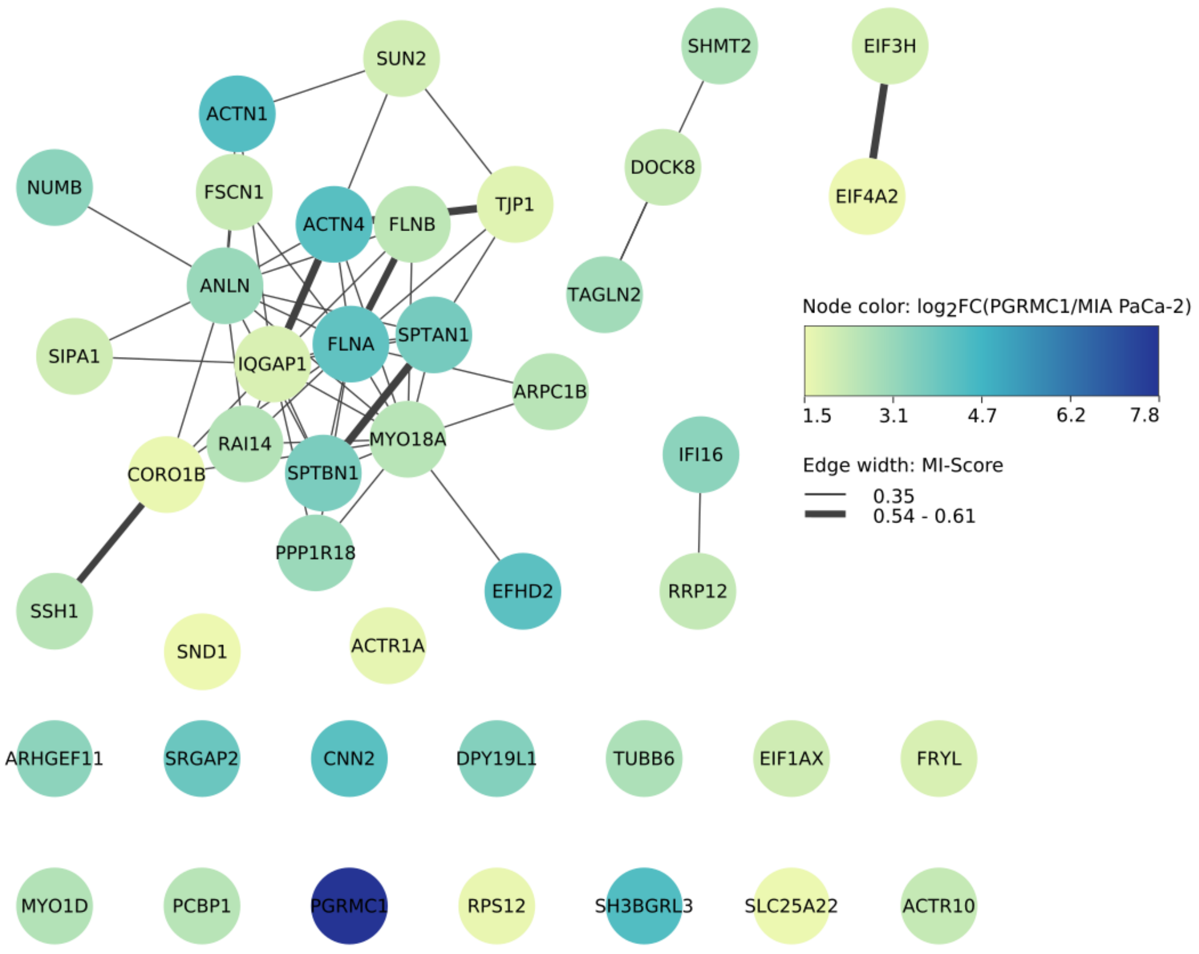

Figure 36: The final network visualization.

##### 7.3.2 Networks generated from multiple group comparisons

This example was produced with the example dataset, using the "Analyze multiple comparisons" feature for the comparisons `PGRMC1__vs__MIAPACA` and `PGRMC1_AG205__vs__MIAPACA_AG205` (we consider enriched proteins, with log2FC $\geq$ 1.5 and adj.p-value $\leq$ 0.05 in at least one of the group comparisons).

The attributes in the gml output from a multi-group comparison are different from those of the single group comparison. Networks from multiple group comparisons have two types of edges:

1. Edges connecting the group comparison with significant proteins.

2. Edges from the IntAct database connecting proteins with proteins.

To distinguish between these types we could set a discrete mapping for the edge color (Stroke Color (unselected)) property in the Edge Style page (Fig. 37b). We can show the different interaction scores of the PPIs by creating a continuous or discrete mapping for the value column.

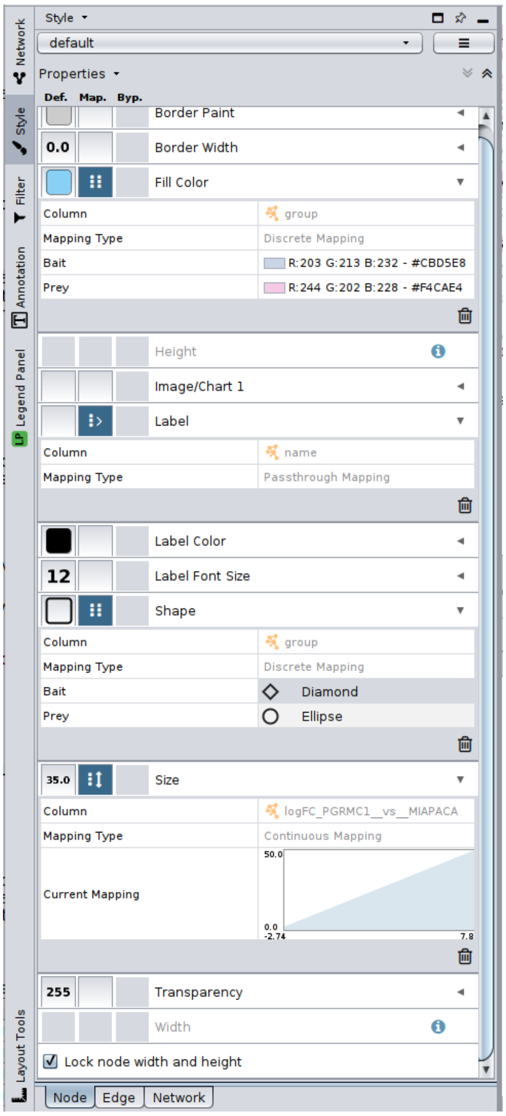

(a) Node style.

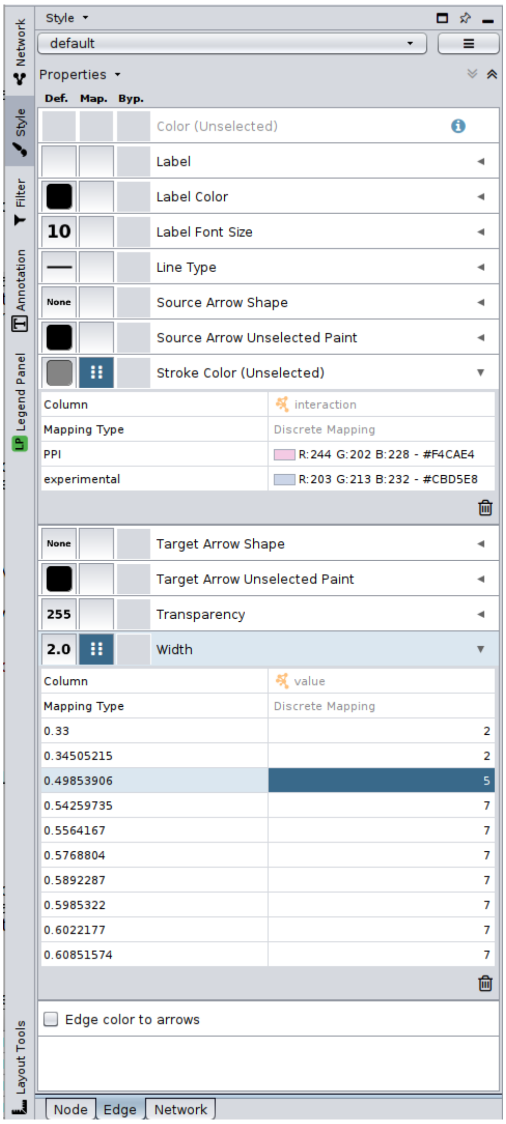

(b) Edge style.

Figure 37: Selected mappings in Cytoscape.

If we want to integrate quantitative information we have to include amicas data table which can be downloaded as csv file in the differential abundance tab. In Cytoscape we have to select:

```
File > Import > Table from File
```

Select the correct parameters (Fig. 38) for the target table (choose the column with the name `key` as `key`) and network collection and fold changes and p-values should now be available in your Node table.

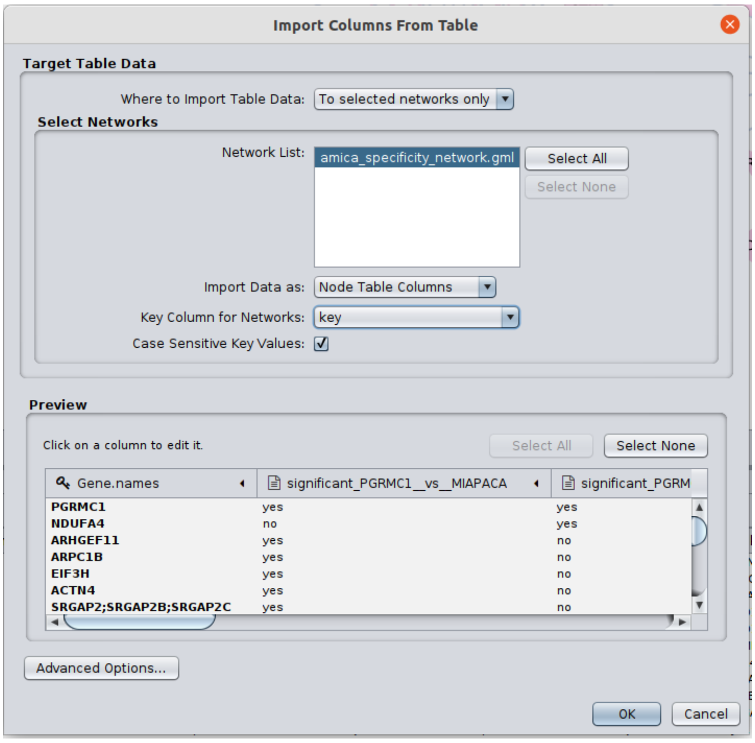

Figure 38: File import and merge options. The `Gene.names` column in the data table corresponds to the `key` column in the gml file.

From here on we can select mappings for the node style. Networks from multiple group comparisons contain two type of nodes in the `group` attribute of the gml file:

1. Bait: a group comparison (in this case `PGRMC1__vs__MIAPACA` and `PGRMC1_AG205__vs__MIAPACA_AG205`).

2. Prey: significant proteins of a group comparison.

We can select a discrete mapping of the group column for the fill color, a passthrough mapping of the name column for the label, a discrete mapping of the group column for the shape and a continous mapping of the logFC_PGRMC1__vs__MIAPACA column for the size, see Fig. 37a).

After applying all these mappings we can export the network visualization, which should look similar like the one shown in Figure 39 (The legend was created in inkscape):

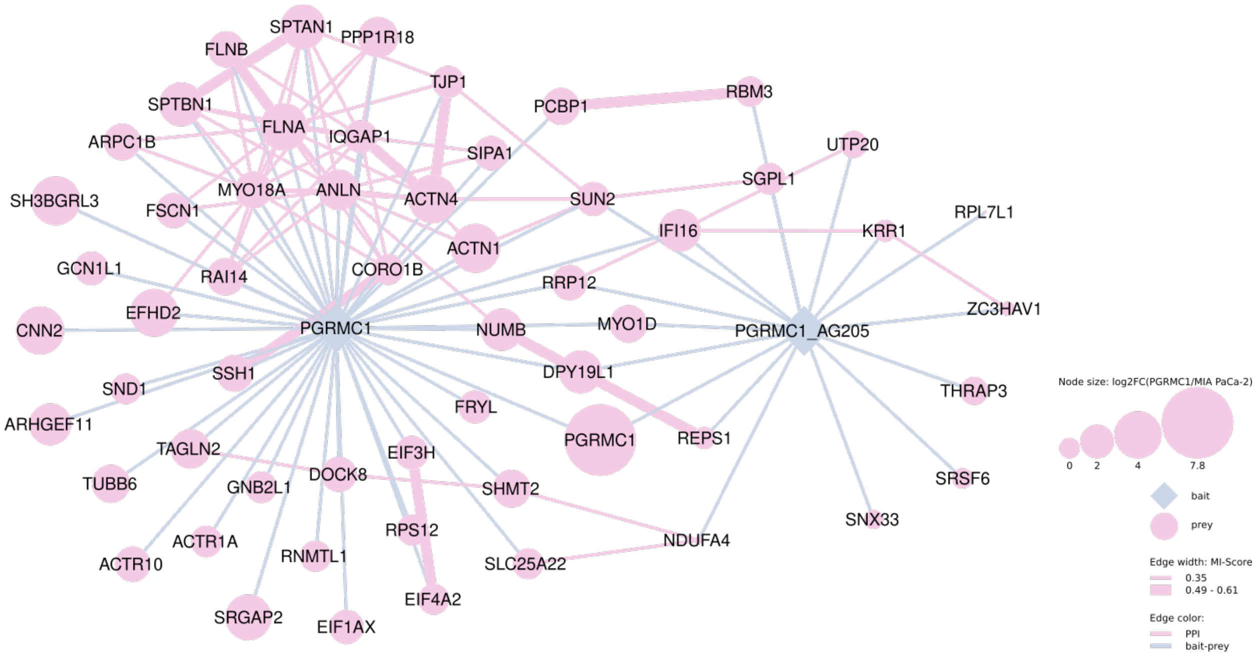

Figure 39: The final network visualization.

From there, you can follow these useful tutorials to visualize the data the way you want:
`https://cytoscape.org/cytoscape-tutorials/protocols/basic-data-visualization`
`https://cytoscape.org/cytoscape-tutorials/protocols/AP-MS-network-analysis`

### Chapter 8

### Comparison with other R-Shiny apps for proteomics data analysis

While other tools such as ProVision, LFQ-Analyst and Eatomics [17, 18, 19] are limited to analyzing MaxQuant's output, amica also automatically recognizes the output of FragPipe, a recent but increasingly popular software platform for the analysis of proteomics data. Additionally, amica allows the upload of any custom tab-separated file, provided that a specification file is uploaded that maps relevant database search tool specific columns to a standard format.

The results of amica's analysis can be downloaded in a specially developed tab-separated format which incidentally is also an accepted input format, facilitating the re-inspection of a dataset at a later timepoint, without having to repeat any of the analysis steps. Furthermore, this allows researchers to share results with collaborators.

Moreover, the ability of converting any analyzed dataset (e.g RNA-Seq data) into the defined amica input format allows for the exploration of data originating from different sources not limited to MS-based proteomics. Combined with amica's unique feature to upload a second file to be compared to the current data input, this enables multi-omics integration.

Furthermore, none of the above mentioned tools are capable of analyzing pilot experiments without replicates. This feature might be useful to users and facilities to decide whether it makes sense to invest resources into a large experiment.

Lastly, amica allows for the integration of quantitative information as well as subcellular localization predictions onto PPI networks. Beside the interactive network displayed in amica, all necessary information needed for visualization in a network visualization tool such as Cytoscape can be downloaded in gml format.

[9] Erich Neuwirth and R Color Brewer. Colorbrewer palettes. *R package version*, 1, 2014.

[15] Uku Raudvere, Liis Kolberg, Ivan Kuzmin, Tambet Arak, Priit Adler, Hedi Peterson, and Jaak Vilo. g: Profiler: a web server for functional enrichment analysis and conversions of gene lists (2019 update). *Nucleic acids research*, 47(W1):W191–W198, 2019.

[16] Paul Shannon, Andrew Markiel, Owen Ozier, Nitin S Baliga, Jonathan T Wang, Daniel Ramage, Nada Amin, Benno Schwikowski, and Trey Ideker. Cytoscape: a software environment for integrated models of biomolecular interaction networks. *Genome research*, 13(11):2498–2504, 2003.

[17] James Luke Gallant, Tiaan Heunis, Samantha Leigh Sampson, and Wilbert Bitter. Provision: a web-based platform for rapid analysis of proteomics data processed by maxquant. *Bioinformatics*, 36(19):4965–4967, 2020.

[18] Anup D Shah, Robert JA Goode, Cheng Huang, David R Powell, and Ralf B Schittenhelm. Lfq-analyst: an easy-to-use interactive web platform to analyze and visualize label-free proteomics data preprocessed with maxquant. *Journal of proteome research*, 19(1):204–211, 2019.

[19] Milena Kraus, Mariet Mathew Stephen, and Matthieu-P Schapranow. Eatomics: Shiny exploration of quantitative proteomics data. *Journal of Proteome Research*, 20(1):1070–1078, 2020.
